# Structural basis for ion selectivity in potassium-selective channelrhodopsins

**DOI:** 10.1101/2022.10.30.514430

**Authors:** Seiya Tajima, Yoon Seok Kim, Masahiro Fukuda, Eamon F.X. Byrne, Peter Y. Wang, Joseph M. Paggi, Koichiro E. Kishi, Charu Ramakrishnan, Syunki Takaramoto, Takashi Nagata, Masae Konno, Masahiro Sugiura, Kota Katayama, Toshiki E. Matsui, Keitaro Yamashita, Hisako Ikeda, Masatoshi Inoue, Hideki Kandori, Ron O. Dror, Keiichi Inoue, Karl Deisseroth, Hideaki E. Kato

**Author notes:** Correspondence (K.D.), (H.E.K.). These authors contributed equally (Seiya Tajima, Yoon Seok Kim).

## Abstract

The KCR channelrhodopsins are recently-discovered light-gated ion channels with high K^+^ selectivity, a property that has attracted broad attention among biologists– due to intense interest in creating novel inhibitory tools for optogenetics leveraging this K^+^ selectivity, and due to the mystery of how this selectivity is achieved in the first place. Indeed, the molecular and structural mechanism for K^+^ selectivity in KCRs has remained especially puzzling since these 7-transmembrane retinal-binding proteins completely lack structural similarity with known K^+^ channels, which generally coordinate K^+^ in a precisely symmetric conduction pathway formed by a tight interface among multiple small monomeric channel subunits (presumably not an accessible mechanism for the large KCR rhodopsin proteins). Here we present the cryo-electron microscopy structures of two KCRs from *Hyphochytrium catenoides* with distinct spectral properties for light absorption and channel actuation, *Hc*KCR1, and *Hc*KCR2, at resolutions of 2.6 and 2.5 Å, respectively. Structural comparison revealed first an unusually-shaped retinal binding pocket which induces rotation of the retinal in *Hc*KCR2, explaining the large spectral difference between *Hc*KCR1 and 2. Next, our combined structural, electrophysiological, computational, and spectroscopic analyses revealed a new solution to the challenging problem of K^+^-selective transport. KCRs indeed do not exhibit the canonical tetrameric K^+^ selectivity filter that specifically coordinates dehydrated K^+^; instead, single KCR monomers form a size exclusion filter using aromatic residues at the extracellular side of the pore which inhibits passage of bulky hydrated ions. This unique feature allows KCRs to function as K^+^ channels under relevant physiological conditions, providing not only a novel mechanism for achieving high K^+^ permeability ratios in biological ion channels, but also a framework for designing the next generation of inhibitory optogenetic tools.

**In Brief:** The first structures of K^+^-selective channelrhodopsins (*Hc*KCR1 and 2) are determined, revealing a K^+^ selectivity mechanism distinctly different from canonical K^+^ channels.

**Highlights:**

- The cryo-EM structures of K^+^-selective channelrhodopsins, *Hc*KCR1 and 2, in nanodisc
- Conditions under which naturally-occurring microbial rhodopsins have a 6-s-*cis* retinal
- Identification of key residues for high K^+^ permeability ratios
- The unique K^+^ selectivity mechanism of KCRs

## INTRODUCTION

Motile organisms typically sense light, one of the most important energy sources and environmental signals, using rhodopsin family proteins. Rhodopsins are largely classified into two groups: microbial (type-1) and animal (type-2) (de Grip and Ganapathy, 2022). Both types contain a seven-helix transmembrane (7TM) domain (opsin) that is covalently bound to a chromophore (retinal) via a Schiff base linkage, but molecular mechanisms for the two types are remarkably different. Most microbial rhodopsins have all-*trans* retinal in the dark state, and light absorption triggers the isomerization from all-*trans* to 13-*cis* configuration. This photoisomerization induces a sequence of structural changes in the opsin (the photocycle), that results in a variety of molecular functions, such as those underpinning ion pumps, ion channels, sensors, and enzymes (Kato, 2021; Nagata and Inoue, 2021). When combined with precise light and viral delivery methods, heterologous expression of these proteins (especially ion channel- and pump-type rhodopsins), enables control of the membrane potential of specific cells in behaving organisms with high spatiotemporal resolution. This experimental approach (optogenetics) has been applied to study the function of neural circuits, analyze the physiology of non-neuronal systems, and treat human diseases (Deisseroth, 2015; Deisseroth and Hegemann, 2017; Emiliani et al., 2022; Sahel et al., 2021).

Non-selective cation channelrhodopsins (cation ChRs or CCRs) from chlorophyte algae, which conduct a wide range of monovalent and divalent cations (e.g. H^+^, Na^+^, K^+^, Ca^2+^), were first applied to enable optogenetics (Deisseroth, 2015) by exciting neurons with light-activated inward currents (Deisseroth and Hegemann, 2017; Bi et al., 2006; Boyden et al., 2005; Ishizuka et al., 2006; Li et al., 2005; Nagel et al., 2005). Following the discovery of the first ChR in 2002 (Nagel et al., 2002), many ChR variants with unique properties in kinetics, conductance, absorption spectrum, ion selectivity, and light sensitivity have been engineered or isolated from nature, greatly expanding the optogenetics toolbox for neuronal excitation (Emiliani et al., 2022). Nevertheless, in contrast to the rapid advance of excitatory optogenetics, the development of tools for neuronal inhibition has lagged. Light-induced neuronal inhibition was first achieved by inward Cl^-^ pumps and outward H^+^ pumps (Chow et al., 2010; Zhang et al., 2007), and more potent inhibition was later achieved by designed and natural Cl^-^-conducting anion channelrhodopsins (anion ChRs or ACRs) (Berndt et al., 2014; Govorunova et al., 2015; Wietek et al., 2014). These ACRs are now used in a wide variety of model organisms including mice, fish, and worms (Antinucci et al., 2020; Berndt et al., 2016; Kato et al., 2018; Mahn et al., 2018; Mohammad et al., 2017). However, variations in Cl^-^ concentration gradients among different subcellular compartments or different developmental stages sometimes causes neuronal excitation from ACR activation, rather than the desired inhibition, thereby limiting the application of these tools (Mahn et al., 2016; Wiegert et al., 2017). Under physiological conditions, repolarization of neuronal membranes universally occurs via an efflux of K^+^ ions. Therefore, K^+^-selective ChRs were long considered tools of interest for neuronal silencing (Wiegert et al., 2017), if they could be created or designed; however, it was assumed to be extremely challenging to engineer or discover them because previously known K^+^ channels generally exhibit a highly conserved domain organization with no similarity to ChRs (Figure S1). Canonical K^+^ channels assemble as a tetramer and the ion-conducting pore is formed by the tetramer interface; each protomer has a highly conserved TVGYG or related motif lining the pore to form a radially-symmetric selectivity filter that specifically coordinates the K^+^ ions (Gouaux and Mackinnon, 2005) (Figure S1B). In contrast, while ChRs form oligomers (dimers or trimers), the ion-conducting pathway is placed within each monomer and is highly asymmetric (Kato, 2021) (Figure S1C). Given this vast structural chasm separating the two types of protein, it was considered possible that naturally-occurring K^+^-selective ChRs would be hard to find and that engineering efforts to combine the two would prove to be extremely challenging.

After identification of the cation-conducting (CCR) and anion-conducting (ACR) families of channelrhodopsin, a third family was discovered in cryptophyte algae and from marine metagenomic datasets; these are termed pump-like channelrhodopsins (PLCRs) (or bacteriorhodopsin-like channelrhodopsins) because they show greater sequence similarity to archaeal pump-type rhodopsins (chloride and proton pumps: halorhodopsins and bateriorhodopsins, respectively) than to canonical ChRs (Govorunova et al., 2016; Marshel et al., 2019; Yamauchi et al., 2017). While PLCRs show relatively high sequence similarity with pump-type rhodopsins, these proteins actually work as ion channels. For example, ChRmine (a recently discovered PLCR; Marshel et al., 2019) exhibits extremely high photocurrents (currently one of the most potent excitatory optogenetic tools; Marshel et al., 2019; Vogt, 2022) and high light sensitivity, as well as a red-shifted action spectrum (Marshel et al., 2019). Interestingly, some PLCRs, such as ChRmine and CCR4 from *Guillardia theta* (*Gt*CCR4), do not permeate divalent cations and display high selectivity for monovalent ions. However, these still transport both Na^+^ and K^+^ (Kishi et al., 2022; Shigemura et al., 2019).

Two microbial rhodopsins isolated from the hyphochytrid protist *Hyphochytrium catenoides* were identified as naturally-occurring light-gated K^+^-selective channels (Govorunova et al., 2022). These new rhodopsins, *Hyphochytrium catenoides* Kalium channelrhodopsins 1 and 2 (*Hc*KCR1 and *Hc*KCR2), are homologous by sequence to previously discovered PLCRs (Kishi et al., 2022; Sineshchekov et al., 2017), but exhibit higher selectivity for K^+^; the K^+^/Na^+^ permeability ratio (*P*_K_/*P*_Na_) of *Hc*KCR1 and 2 reaches to ∼23 and ∼17, respectively, much greater than those of other PLCRs (e.g. *P*_K_/*P*_Na_ of both *Gt*CCR4 and ChRmine are ∼0.9) (Shigemura et al., 2019; data not shown), canonical ChRs (e.g. *P*_K_/*P*_Na_ of ChR2 from *Chlamydomonas reinhardtii* (*Cr*ChR2) is ∼0.5) (Nagel et al., 2003), or even some canonical K^+^ channels (e.g. *P*_K_/*P*_Na_ of KcsA and mouse SLO3 are ∼11 and 5-10, respectively) (Meuser et al., 1999; Santi et al., 2009).

The potential for KCRs as potent inhibitory optogenetic tools was demonstrated in mammalian neurons (Govorunova et al., 2022). However, the fundamental question of how K^+^ selectivity is achieved in channelrhodopsins remains to be answered. Understanding the structural basis of K^+^ selectivity by KCRs would be enormously valuable, not only as a new paradigm for understanding how ion channel proteins can achieve K^+^ selectivity, but also to provide a framework for the creation of next-generation KCR-based optogenetic tools. Here, we present the cryo-electron microscopy (cryo-EM) structures of *Hc*KCR1 and 2, at resolutions of 2.6 and 2.5 Å, respectively. The structural information, along with spectroscopic, electrophysiological, and computational analyses, reveals the unique mechanisms of initial photoreactions, color tuning, and high K^+^ permeability ratios of KCRs.

## RESULTS

### Overall Structural Comparison between *Hc*KCR1, *Hc*KCR2, and ChRmine

For structural determination, we expressed *Hc*KCR1 and 2 (residues 1-265 for both) in Sf9 insect cells, and reconstituted the purified proteins into lipid nanodiscs formed by the scaffold protein MSP1E3D1 and soybean lipids (STAR methods). Using cryo-EM, we solved the structures of the *Hc*KCR1 and 2 in the dark state to overall resolutions of 2.6 Å and 2.5 Å, respectively (Figures S2A-W; Table S1). The high-resolution density maps allowed us to accurately model the vast majority of both *Hc*KCRs (residues 6-260 for *Hc*KCR1 and 2-260 for *Hc*KCR2), as well as water molecules, lipids, and the all-*trans* retinal whose conformer was also validated by high-performance liquid chromatography (HPLC) analysis (Figures S2O-W, S3A, and S3B). The N-terminal residue of *Hc*KCR2, P2, is surrounded by four residues (P95, F96, W100, and Y101) in the structure and there is no space for the first methionine (Figure S2V). This is consistent with previous findings revealing that the first methionine is post-translationally cleaved off when the second and third residues are proline and non-proline, respectively (the third residue is phenylalanine in *Hc*KCR2) (Wingfield, 2017).

Both *Hc*KCR1 and 2 form a trimer (Figures 1A and 1B), as was also observed in ChRmine, the only PLCR for which high-resolution structural information is available (Kishi et al., 2022; Tucker et al., 2022). The trimerization is mainly achieved by the direct and lipid-mediated interactions among transmembrane helices (TMs) 1-2 and TMs 4-5 of adjacent protomer, and the center of the trimer interface is filled with six lipid molecules (Figures 1A and 1B). The monomer of *Hc*KCR1 and 2 consists of an extracellular N-terminal region (residues 6-21 for *Hc*KCR1 and 2-21 for *Hc*KCR2), an intracellular C-terminal region (residues 255-260 for both), and 7-TM domains (within residues 22-254 for both), connected by three intracellular loops (ICL1-3) and three extracellular loops (ECL1-3) (Figures 1C and 1D). The overall structures of *Hc*KCR1 and 2 are almost identical with a Cα root-mean-square deviation (r.m.s.d.) of only 0.51 Å and only minor differences in the N-terminal region, ICLs, and ECLs (Figure 1E).

**Figure 1.**
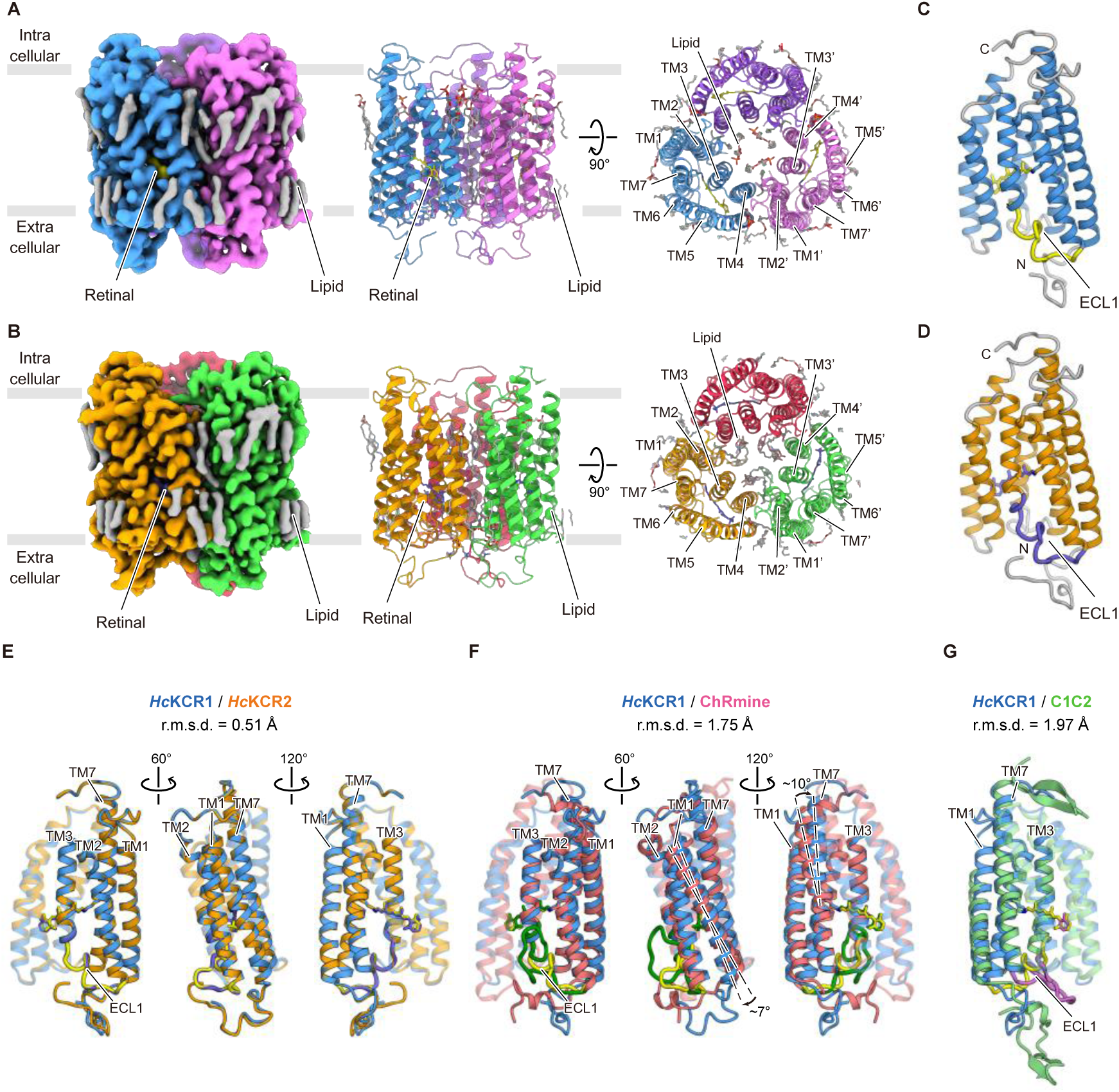
Cryo-EM structures of *Hc*KCR1 and *Hc*KCR2. (A) Cryo-EM density map (left) and ribbon representation of the *Hc*KCR1 homotrimer viewed parallel to the membrane (middle) and viewed from the intracellular side (right), colored by protomer (blue, magenta, and purple), retinal (yellow), and lipid (grey), respectively. Grey bars indicate approximate location of the lipid bilayer. (B) Cryo-EM density map (left) and ribbon representation of *Hc*KCR2 homotrimer viewed parallel to the membrane (middle) and viewed from the intracellular side (right), colored by protomer (orange, green, and red), retinal (purple), and lipid (grey), respectively. Grey bars indicate approximate location of the lipid bilayer. (C and D) Monomeric structures of *Hc*KCR1 (C) and *Hc*KCR2 (D). 7-TM domains of *Hc*KCR1 and *Hc*KCR2 are colored in blue and orange, respectively. Retinal and ECL1 are colored in yellow for *Hc*KCR1 and purple for *Hc*KCR2, respectively. (E–G) Structural comparisons among *Hc*KCR1, *Hc*KCR2, ChRmine, and C1C2. *Hc*KCR1 (blue) superimposed onto *Hc*KCR2 (orange) (E), ChRmine (red) (F), and C1C2 (green) (G) from different angles. Retinal and ECL1 are colored in yellow (*Hc*KCR1), purple (*Hc*KCR2), green (ChRmine), and pink (C1C2). TMs 4-6 are displayed with transparency for clarity. Compared to ChRmine, TM1 and the cytoplasmic half of TM7 of *Hc*KCR1 are tilted by about 7 and 10 degrees, respectively.

*Hc*KCRs also superpose well onto ChRmine, but with several structural differences (Cα r.m.s.d. between *Hc*KCR1 and ChRmine is 1.75 Å) (Figure 1F). First, both the N- and C-terminal regions in ChRmine have short α-helices running almost parallel to the membrane, which are absent in the *Hc*KCRs (Figure 1F). Second, except for ECL3, all ICLs and ECLs have significantly different conformations. ECL1 in particular, which distinguishes PLCRs from the rest of the ChR families, is ∼6 residues shorter than ChRmine, and the entire loop is packed more closely to the core of the helix bundle (Figure 1F). Third, TM1 and the C-terminal half of TM7 are tilted about 7 and 10 degrees, respectively, relative to the rest of the helical bundle. The C-terminal TM7 helix is also ∼1.5 turns longer than that of ChRmine (Figure 1F), making it more similar to that of canonical CCRs such as C1C2 (the chimera derived from *Cr*ChR1 and *Cr*ChR2) (Figure 1G). In PLCRs, residues from TM1, 2, 3, 7, and ECL1 form the core of the ion-conducting pathway within each monomer (Kishi et al., 2022; Tucker et al., 2022), so the structural differences of TM1, 7, and ECL1 observed in *Hc*KCRs change the shape of the pathway, to be discussed in more details later.

### The Schiff Base Region

Microbial rhodopsins have an all-*trans* retinal molecule covalently bound to a conserved lysine residue on TM7 via a Schiff base linkage. The Schiff base is protonated in the dark and this positive charge must be stabilized by one or more nearby acidic residues for efficient isomerization of retinal (Tahara et al., 2018). Initial reactions triggered by light absorption include retinal isomerization and subsequent proton transfer from the Schiff base to a nearby acidic residue or water molecule. The residues stabilizing the Schiff base proton and receiving the proton in the photo-intermediate state (M intermediate) have been historically termed the Schiff base counterion(s) and the proton acceptor, respectively (Ernst et al., 2014). The precise architecture of the Schiff base region is closely linked to several key properties of microbial rhodopsins (Ernst et al., 2014), so we next focused on this region.

Our previous study revealed that the Schiff base region of ChRmine is strikingly different from those of other microbial rhodopsins (Kishi et al., 2022); TM3 is unwound in the middle of the membrane, and two aspartates, the strong candidates for the Schiff base counterion and proton acceptor, are placed on TM3/ECL1 and TM7. The first aspartate (D115 in ChRmine) faces away from the Schiff base proton, and the second aspartate (D253 in ChRmine) is fixed by two hydrogen bonds with Y85 on TM2 and Y116 on TM3 (Figure 2A, right). These features were also observed in *Hc*KCR1 and 2, suggesting that the architecture of the Schiff base region is conserved among PLCRs (Figures 2A and S4A). However, there are still several differences between *Hc*KCR1, 2, and ChRmine: K84 points towards the extracellular side in *Hc*KCRs (Figures 2A and S2S); no water molecules are observed between the Schiff base proton and the two aspartates (D105 and D229 in *Hc*KCRs) in *Hc*KCRs (Figure 2A); and the highly conserved arginine residue (Fig. S1A) on ECL1 (R112 in ChRmine) is replaced by a tryptophan residue (W102 in *Hc*KCRs) in *Hc*KCRs (Figures 2A and S1A). These differences motivated us to further characterize the functions of D105 and D229.

**Figure 2.**
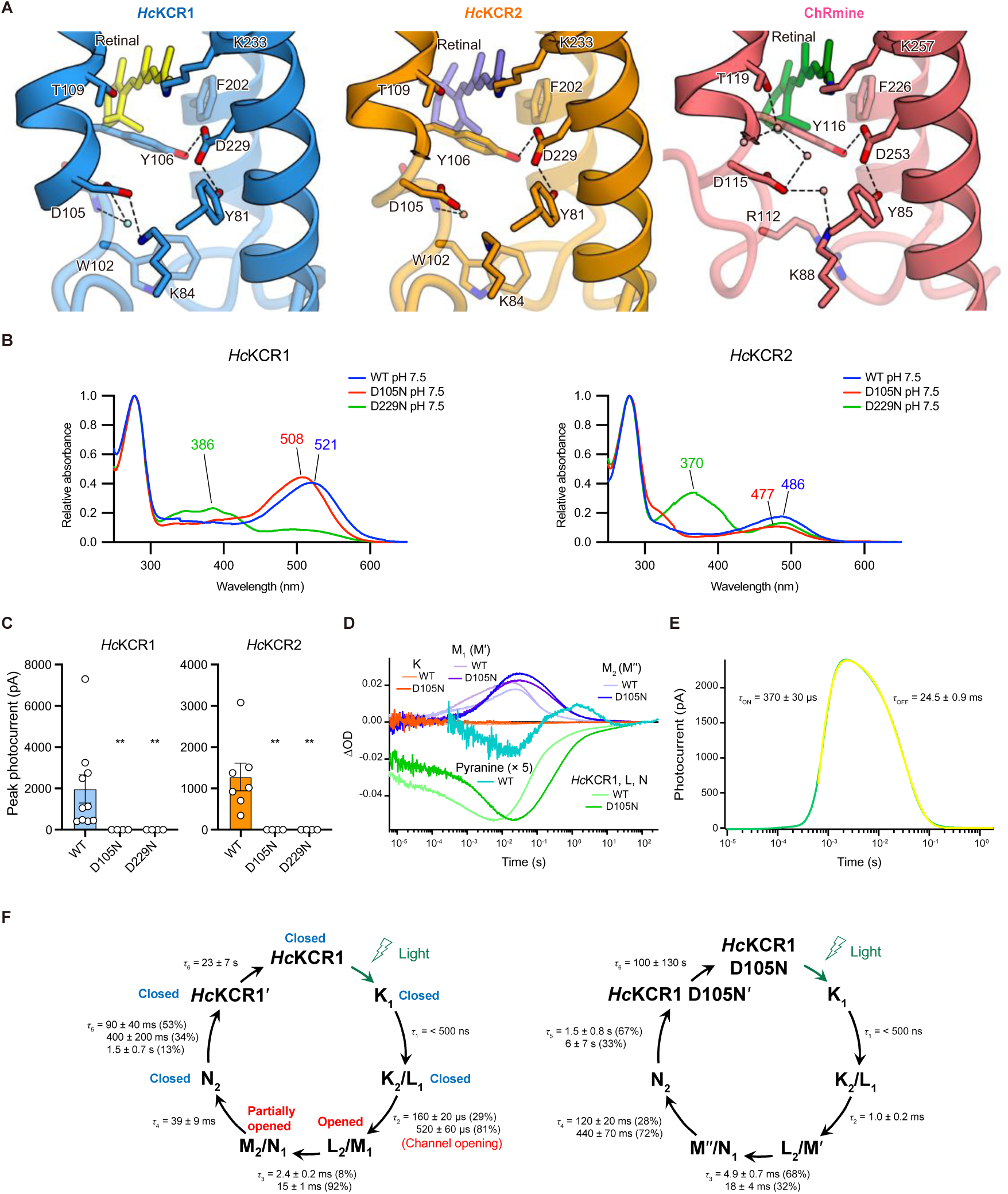
The Schiff base region. (A) The Schiff base regions of *Hc*KCR1 (left), *Hc*KCR2 (middle), and ChRmine (right). Water molecules are represented by spheres. The black dashed lines indicate H-bonds. (B) Absorption spectra of *Hc*KCR1 (left) and *Hc*KCR2 (right) at pH 7.5. The traces of WT, D105N, and D229N are colored in blue, red, and green, respectively. The λ_max_ values are shown above each trace. (C) Photocurrent amplitudes of WT, D105N and, D229N of *Hc*KCR1 (left) and *Hc*KCR2 (right), respectively. Mean ± SEM (n = 4–10); Kruskal-Wallis test with Dunnett’s test. ** p < 0.01. (D) Time-series traces of absorption change for *Hc*KCR1 WT and D105N mutant at specific wavelength. For *Hc*KCR1 WT, the probe wavelength at 617 nm (light red), 480 nm (light green), 384 nm (light purple), and 404 nm (light blue) corresponds to K, L and N, M_1_, and M_2_ intermediates, respectively. The corresponding wavelengths of K, L and N, Mʹ, and Mʹʹ intermediates for *Hc*KCR1 D105N are 609 nm (red), 515 nm (green), 378 nm (purple), and 394 nm (blue), respectively. The cyan line represents the absorption changes of pyranine monitored at 454 nm. (E) Transient photocurrent changes of *Hc*KCR1 induced by pulsed flash laser. Green and yellow lines indicate the raw trace and the fitting curve, respectively. (F) Photocycle schemes of *Hc*KCR1 WT (left) and D105N mutant (right) determined by flash photolysis experiment shown in (D).

First, to assign protonation states of these two aspartates and to identify which aspartate works as the primary counterion, we measured the absorption spectra of wild-type (WT), D105N, and D229N mutants of both *Hc*KCRs (Figures 2B, S3C, and S3D). The λ_max_ of WT, D105N, and D229N mutants of *Hc*KCR1 at neutral pH is 521, 508, and 386 nm, respectively, demonstrating that protonation of D105 causes only a small blue-shift in the absorption spectrum (∼13 nm), while protonation of D229 causes a much larger blue-shift (∼135 nm), which can be explained by concomitant deprotonation of the Schiff base nitrogen (Figure 2B, left). The same trend was also observed in *Hc*KCR2 (Figure 2B, right). This finding suggests that both D105 and D229 are deprotonated in the dark state, but only the deprotonation of D229 is necessary to stabilize the positive charge of the Schiff base proton; in other words, D229 is the primary counterion. This is strikingly different from ChRmine, in which both of the corresponding aspartate residues (D115 and D253) are essential counterions of the Schiff base proton (Kishi et al., 2022).

However, surprisingly, our electrophysiology experiments with these mutants showed that channel function is completely abolished not only for D229N but also for D105N (Figures 2C and S5). To understand the reason, we next performed laser flash photolysis and laser patch clamp experiments (Figures 2D-F, S3E, and S3F). These experiments revealed that *Hc*KCR1 has eight intermediates (K_1_, K_2_, L_1_, L_2_, M_1_, M_2_, N_1_, N_2_) in its photocycle, with M_1_ and M_2_ representing the open state, consistent with a previous study (Govorunova et al., 2022). We further measured the photocycle of the D105N mutant and found that the rise and decay of M intermediate become significantly slower in this mutant (Figures 2D and 2F), suggesting that D105 works as the proton acceptor. This interpretation is also supported by the flash photolysis experiment of WT *Hc*KCR1 in the presence of pyranine, a pH-sensitive dye (Kano and Fendler, 1978), showing that the Schiff base proton is released to the bulk solvent later than the rise of the M_1_ (Figures 2D); this result indicates that the Schiff base proton is not directly released to water but is transferred to an acidic residue in the Schiff base region. Notably, the shapes of the absorption spectra of M_1_ and M_2_ intermediates in the D105N mutant are significantly different from those in WT *Hc*KCR1 (Figure S3F), thus it can be inferred that the structures of D105N mutant in these intermediates, which are newly denoted as M’ and M’’ (Figure 3F), are also different from those of WT, and thereby channel function of this mutant is compromised. Overall, these results suggest that D105 does not work as the Schiff base counterion in the dark state but works as the proton acceptor in the M intermediate, and the proton transfer to the D105 would be an important step for correct channel gating.

**Figure 3.**
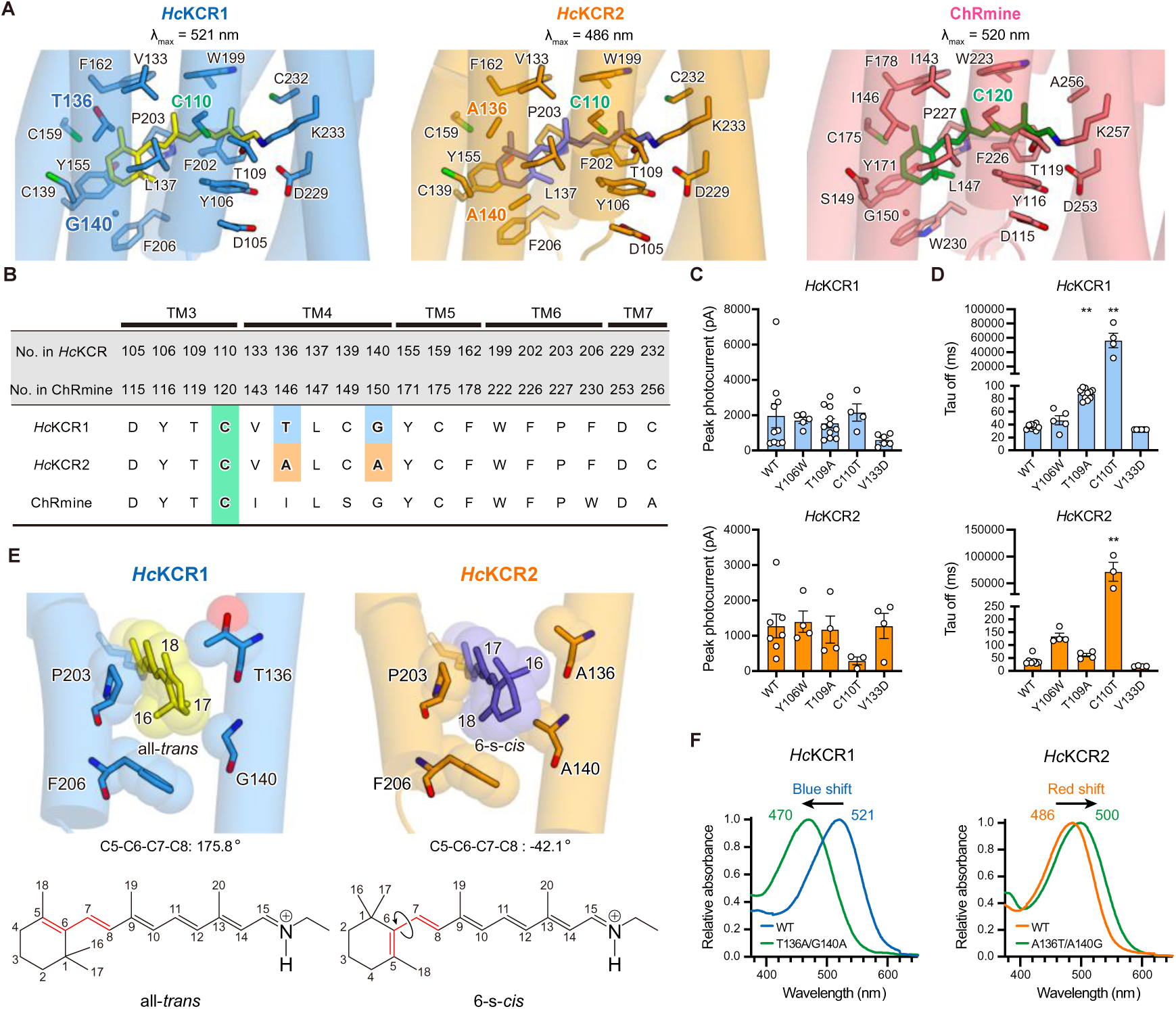
Retinal binding pocket. (A) Retinal binding pockets of *Hc*KCR1 (left), *Hc*KCR2 (middle), and ChRmine (right). Residues forming the retinal binding pockets are shown in stick model form. (B) Sequence alignment for residues in the retinal binding pocket. (C) Peak photocurrent amplitudes of WT and four mutants of *Hc*KCR1 (top) and *Hc*KCR2, respectively (bottom). Mean ± SEM (n = 3–11); Kruskal-Wallis test with Dunnett’s test. (D) τ_off_ of WT and four mutants of *Hc*KCR1 (top) and *Hc*KCR2 (bottom), respectively. Mean ±SEM (n = 3–11); Kruskal-Wallis test with Dunnett’s test. ** p < 0.01. (E) β-ionone rings of *Hc*KCR1 and *Hc*KCR2 (top), and chemical structures of all-*trans* and 6-s-*cis* retinal (bottom). Red lines represent C5-C6-C7-C8 bonds. (F) Absorption spectra of *Hc*KCR1 and 2 WT and their swapping mutants (T136A/G140A for *Hc*KCR1 and A136T/A140G for *Hc*KCR2). The λ_max_ values are shown above each trace.

### The Retinal Binding Pocket

The residues surrounding the retinal chromophore are important determinants of key ChR properties, including kinetics and absorption spectrum (Berndt et al., 2009; Kamiya et al., 2013; Oda et al., 2018), thus we next focused on this region.

The residues comprising the retinal binding pocket are very similar between *Hc*KCR1, 2, and ChRmine (Figure 3A); 12 of 18 residues are conserved between *Hc*KCRs and ChRmine, and only two residues are different between *Hc*KCR1 and 2 (Figure 3B). To understand the function of these residues, we first introduced mutations to Y106 and T109 in *Hc*KCRs (Y116 and T119 in ChRmine) because a previous study showed that Y116W and T119A mutations in ChRmine significantly decelerate and accelerate off kinetics, respectively (Tucker et al., 2022). However, we found that the effects of corresponding mutations in *Hc*KCRs are very different; the Y106W mutation moderately decelerates the off kinetics of only *Hc*KCR2, and the T109A mutation does not accelerate but decelerates the off kinetics of only *Hc*KCR1 (Figures 3C, 3D, S5, S6A, and S6B). Y106 and T109 are part of the retinal binding pocket as well as part of the Schiff base region (Figure 2A), thus the differences in the Schiff base region described above likely account for the differences in mutational effects among *Hc*KCR1, 2, and ChRmine.

Next, we introduced mutations to C110 and V133, for which threonine, serine, or alanine mutants are known to significantly prolong off-kinetics in *Cr*ChR2 (Berndt et al., 2009; Bamann et al., 2010; Yizhar et al., 2011), giving rise to the step-function opsins (SFOs) which have found broad optogenetic application in neuroscience. Although a previous attempt to transfer this mutation to PLCRs was not successful (Sineshchekov et al., 2020), the result was strikingly successful here for both *Hc*KCR1 and 2 (Figures 3C and 3D). The C110T mutation increased the τ_off_ of *Hc*KCR1 and 2 by ∼1500- and ∼1800-fold, respectively (Figure 3D); notably, the *Hc*KCR1 C110T mutant still shows comparable channel activity to WT (Figure 3C). As far as we know, this is the first study to create a step-function opsin in the PLCR family, and the *Hc*KCR1 C110T mutant is expected to work as a powerful optogenetic tool for long-timescale inhibition.

*Hc*KCR1 and 2 show different spectral properties; λ_max_ of *Hc*KCR1 and 2 is 521 nm and 486 nm, respectively (Figure 2B). The retinal binding pockets of *Hc*KCR1 and 2 are very similar, with the only differences at positions 136 and 140 near the β-ionone ring of the retinal (Figures 3A and 3B), providing an excellent opportunity to test spectral mechanisms. In all reported structures of naturally-occurring microbial rhodopsins, the retinal has a 6-s-*trans* form in the binding pocket (Figure S4B). However, during the structural refinement of *Hc*KCR2 (STAR Methods), we noticed that the 6-s-*trans* retinal exhibits a significant steric clash between the C_17_ atom of the retinal and the methyl group of A140, and strong extra density was observed next to the C_18_ atom (Figure S2X, top). This suggests that the β-ionone ring should be rotated in the *Hc*KCR2 structure, and surprisingly, when we modeled 6-s-*cis* retinal, this conformation perfectly fits the density (Figures S2X, bottom). This result indicates that these two residues (A136 and A140 in *Hc*KCR2) create a steric clash with the C_17_ atom and simultaneously make space to accommodate the C_16_ atom, to induce the rotation of the β-ionone ring (Figure 3E). The ∼220 degrees rotation of the ring shrinks the π-conjugated system of retinal and thereby induces a ∼35 nm spectral shift (Figures 2B and 3E). This is in good agreement with a previous study that showed a designed ChR with glycine and alanine at the same positions, C1C2GA, exhibits retinal rotated by ∼210 degrees and a spectrum blue-shifted by ∼20 nm (Figure S4B) (Kato et al., 2015a). To further test our hypothesis, we swapped these two residues between *Hc*KCR1 and 2 and confirmed that the T136A/G140A mutation to *Hc*KCR1 and A136T/A140G mutation to *Hc*KCR2 cause the predicted blue- and red-shifts, respectively (Figure 3F). The impact of these two residues determining the orientation of the β-ionone ring in the binding pocket largely explains the spectral difference between *Hc*KCR1 and 2. To our knowledge, *Hc*KCR2 is the first naturally-occurring microbial rhodopsin for which a 6-s-*cis* configuration of the retinal has been experimentally demonstrated.

### Ion-conducting Pore and K^+^ Selectivity

The three major classes of ChRs including the PLCRs (Kato et al., 2012, 2018; Kim et al, 2018; Kishi et al., 2022), although assembling as multimers, each possess an ion-conducting pore within the monomer, formed by TM1, 2, 3, and 7. For example, the PLCR ChRmine was discovered to form a trimer with a large opening in the middle of the trimer; although mutations in this region can modulate ion selectivity (Kishi et al., 2022), this opening was not predicted or shown to form a conducting pore for ChRmine in Kishi et al. (2022). In the dark state, the monomer pore is divided into the intracellular and extracellular vestibules (IV and EV) by two or three constriction sites, which are called intracellular, central, and extracellular constriction sites (ICS, CCS, and ECS) (Figure S4C) (Kato, 2021).

*Hc*KCRs have a relatively similar sequence to archaeal pump-type rhodopsins (Figure S1A), but with larger cavities due to structural differences of the pore-forming helices (Figures 4A and S4C). Notably, due to the unwinding of TM3 in the middle of the membrane, not only TM1, 2, 3, and 7, but also ECL1, significantly contribute to the creation of the EV, as observed in the ChRmine structure (Figures 1F and 4A) (Kishi et al., 2022). The overall location of the cavities in *Hc*KCRs is very similar to ChRmine but three notable differences are observed between them.

**Figure 4.**
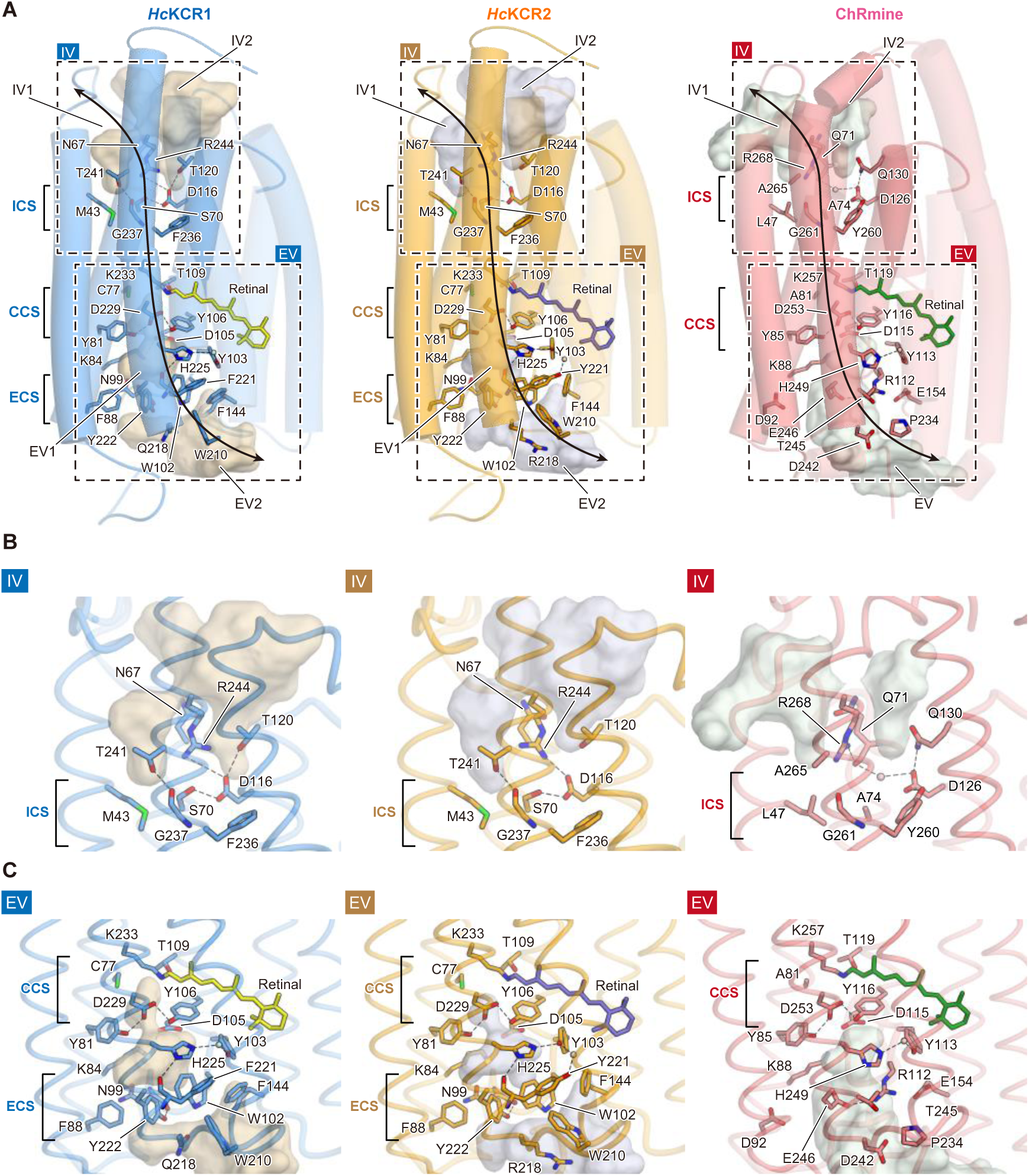
Ion-conducting cavities. (A) Comparison of ion-conducting cavities between *Hc*KCR1 (left), *Hc*KCR2 (middle), and ChRmine (right). TMs 4-6 are displayed with higher transparency. The residues located along the cavities are shown in stick model form. Intra- and extracellular cavities are calculated with the program HOLLOW. The black dashed rectangles indicate the IV and EV regions highlighted in (B) and (C), respectively. The black dashed lines and arrows represent H-bonds and the putative ion-conducting pathway, respectively. Locations of ICS, CCS, and ECS are indicated at the left side of each panel. (B-C) IV (B) and EV (C) of *Hc*KCR1 (left), *Hc*KCR2 (middle), and ChRmine (right). Cavities are calculated with the program HOLLOW, and the black dashed lines indicate H-bonds. Locations of ICS, CCS, and ECS are indicated at the left side of each panel.

First, both *Hc*KCRs and ChRmine exhibit two IVs (IV1 and IV2) divided by a conserved arginine on TM7 (R244 in *Hc*KCRs and R268 in ChRmine), and they are occluded by the ICS, but the interaction network in ICS is significantly different. In ChRmine, R268 forms the H-bond with Q71, and D126 has direct H-bond interactions with both Q130 and Y260 and water-mediated H-bond interaction with Q71. This H-bond network, together with L47, A74, and G261, makes the ICS (Figure 4B, right). In contrast, in *Hc*KCRs, R244 approaches TM3 because of the ∼10 degrees tilt of the cytoplasmic half of TM7 (Figure 1F) and forms a salt bridge with D116 (D126 in ChRmine). Moreover, A74, Q130, and Y260 in ChRmine are replaced by S70, T120, and F236, respectively, resulting in the significant rearrangement of the H-bond network centered on D116 (Figure 4B, left and middle).

Second, the EV in *Hc*KCRs extends deeper into the core of the bundle than ChRmine and indeed reaches the Schiff base (Figure 4C). While the architecture of the Schiff base region is similar between *Hc*KCRs and ChRmine (Figure 2A), the small conformational difference of K84 enlarges the pore in *Hc*KCRs, and the EV extends close to the Schiff base-forming lysine (K233), as observed in *Gt*ACR1 (Figure S4C). As a result, not only the counterion complexes (D155, D229, Y81, and Y106) but also C77, T109, and K233 contribute to defining the CCS in *Hc*KCRs (Figure 4C, left and middle).

Finally, and most importantly, the shape and the surface property of *Hc*KCRs’ EV are strikingly different from that of ChRmine (Figure 4C, left and middle). Several hydrophilic residues that line the EV in ChRmine, including D92, R112, E154, T245, and E246 are replaced by aromatic residues in *Hc*KCRs (F88, W102, F144, F/Y221, and Y222), and they make the EV’s surface more hydrophobic (Figure 4C, left and middle). Moreover, W102 and Y222 protrude to the center of EV and make a new constriction (ECS) with ECL1. As described earlier, *Hc*KCRs’ ECL1 is positioned closer to the core of the helix bundle compared to that of ChRmine (Figure 1F), and allows N99 on ECL1 to forms a H-bond with Y222 and thereby separate the EV into two cavities (Figure 4C, left and middle). The replacement of arginine with tryptophan (W102) in *Hc*KCRs also causes the rotameric change of histidine (H225 in *Hc*KCRs) and generates a new H-bond between H225 and F/Y221 (Figure 4C, left and middle). Overall, these aromatic residues create unique EVs whose shape and properties are different from other microbial rhodopsins (Figures S1A and S4C).

To understand the mechanism of K^+^ selectivity by KCRs, we introduced mutations to the residues placed along the IVs and EVs of *Hc*KCR1 and measured their reversal potentials (E_rev_) (Figures 5A, 5B, S5, S6A, and S6B). WT *Hc*KCR1 exhibits E_rev_ of −68.4 ± 1.3 mV and a permeability ratio (*P*_K_/*P*_Na_) of 25.7 ± 2.7, consistent with a previous study (Govorunova et al., 2022), indicating function as a K^+^-selective channel with minor Na^+^ conductance. We found that most mutations had negligible effects on selectivity, but the mutations to W102, D116, F221, Y222, and H225 caused significant changes in E_rev_ (Figures 5A, 5B, S5, S6A, and S6B). Strikingly, we found that mutants F221A, H225F, H225A, and H225Y became *more* selective to K^+^, with significantly hyperpolarized E_rev_ (−82.1 ± 8.1, −82.0 ± 1.9, −84.6 ± 4.0, and −85.4 ± 3.5 mV, respectively) (Figure 5B). By contrast, W102Q and Y222A mutants became almost non-selective to Na^+^ vs. K^+^ with strikingly depolarized E_rev_ (−6.25 ± 1.5 and −10.6 ± 1.5 mV, respectively). A more conservative mutation at the Y222 position (Y222F) caused an intermediate depolarization in E_rev_ (−44.2 ± 2.9 mV), suggesting that high K^+^ selectivity permeability ratio depends upon bulkiness of the side chains comprising the EV. This finding also agrees well with the previously reported data indicating that KCR selectivity is inversely proportional to the size of hydrated substrate cations (Govorunova et al., 2022; Zhong et al., 2015). These four bulky aromatic residues are localized in the EV, suggesting that W102, F221, Y222, and H225 could assemble and form an ion selectivity filter in *Hc*KCR1 (Figure 5A). This idea is further supported by the homology models of Y222A and W102Q mutants created from the WT cryo-EM structure; the packing interactions between these four residues become weaker in both mutants, and especially Y222A mutant shows a strikingly enlarged cavity compared to WT (Figures 5C and 5D). The disruption of the constriction at the filter region would cause the loss of K^+^ selectivity in these mutants.

**Figure 5.**
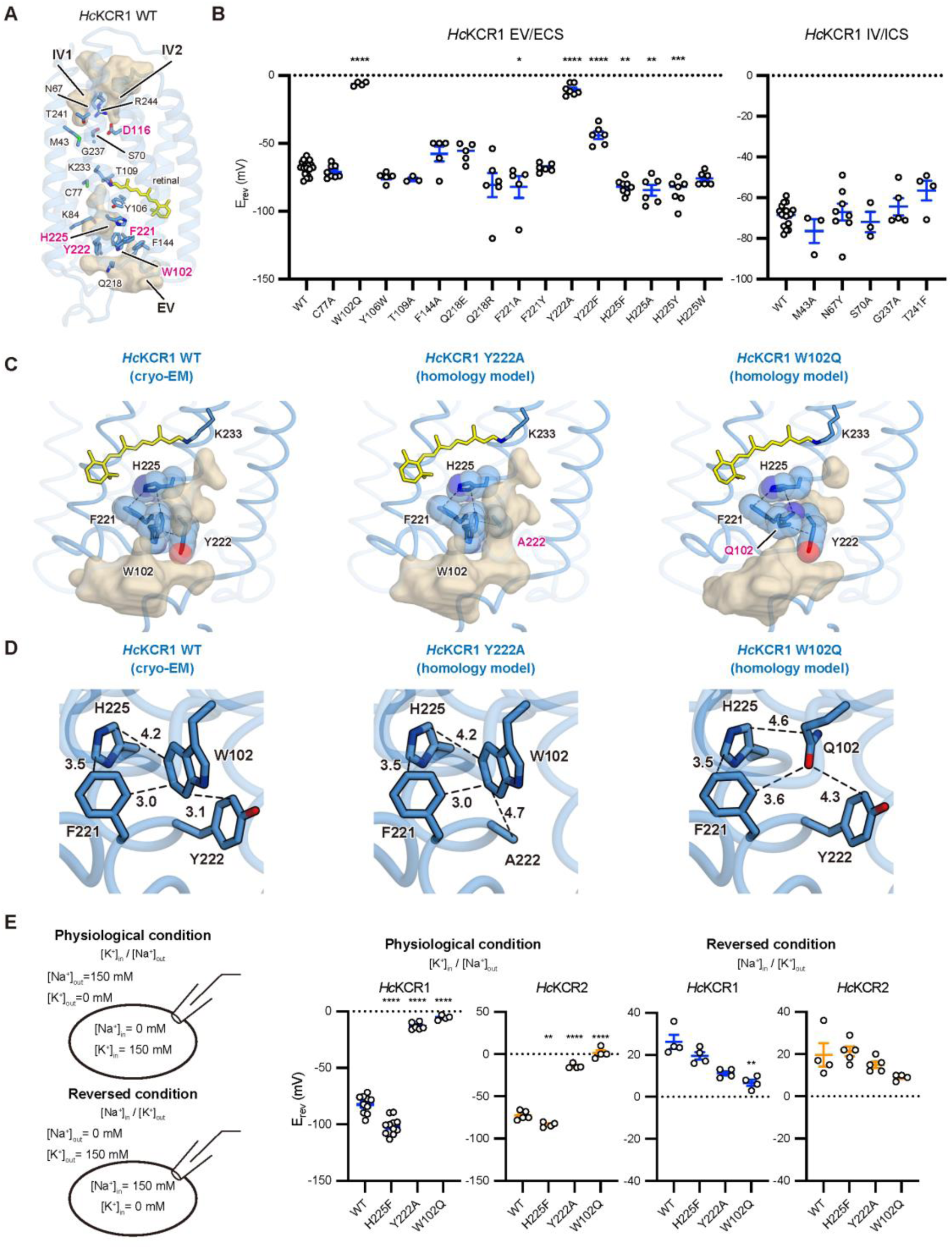
K^+^ selectivity filter. (A, B) Patch clamp characterization of *Hc*KCR1 and *Hc*KCR2. (A) Residues along the ion-conducting cavities in *Hc*KCR1 WT. Residues where mutation significantly changes E_rev_ are highlighted in magenta. (B) E_rev_ summary for mutations of the residues shown in (A). Mutants are categorized as the mutants of EV or ECS (left) and IV or ICS (right). Mean ± SEM (n = 3–17); one-way ANOVA with Dunnett’s test. * p < 0.05 ** p < 0.01, *** p < 0.001, **** p < 0.0001. (C, D) The selectivity filter region of *Hc*KCR1 WT (cryo-EM structure, left), Y222A mutant (homology model, middle), and W102Q mutant (homology model, right), viewed parallel to the membrane (C) and viewed from the extracellular side (D). Cavities are calculated with the program HOLLOW, and the black dashed lines indicate the closest distance between atoms of adjacent amino acids. (E) Patch clamp characterization of *Hc*KCR1 and *Hc*KCR2 under physiological and reversed ion balance conditions. At left: ion concentrations for voltage-clamp recordings. E_rev_ summary for WT and three mutants of *Hc*KCR1 and *Hc*KCR2 under physiological (middle) or reversed (right) conditions. Mean ± SEM (n = 4–21); one-way ANOVA with Dunnett’s test. ** p < 0.01 **** p < 0.0001.

Do these residues select for K^+^? The EV, where these residues are located, is actually the exit site for the substrate K^+^ under typical ion balance conditions, as under physiological electrochemical gradients, K^+^ flows preferentially from the intracellular to the extracellular side (while Na^+^ flows in the opposite direction). Since the EV will predominantly serve as the exit site for K^+^ as well as the entry site for Na^+^ under physiological conditions, we considered that *Hc*KCR1 might achieve its high K^+^ permability ratio under typical chemical gradients (high K^+^/low Na^+^ intracellularly and low K^+^/high Na^+^ extracellularly) chiefly by preventing entry of Na^+^ from the extracellular side via these aromatic amino acids. With robust outward K^+^ currents flowing, little inward Na^+^ would be expected to flow (especially if an aromatic size-exclusion filter deterred flux of the larger hydrated Na^+^ ion in the presence of high flux of the smaller hydrated K^+^ ion). But inward Na^+^ currents would still be possible in the absence of competing K^+^ ions from outside the cell, if at strongly negative membrane potentials such that intracellular K^+^ ions were no longer flowing outward down an electrochemical gradient.

A prediction of this hypothesis would be that robust Na^+^ currents would be observed through KCRs under altered electrochemical gradient conditions: either in the form of 1) inward Na^+^ currents at membrane potentials more negative than the reversal potential for smaller (hydrated) K^+^ ions that could otherwise flow outward and outcompete larger hydrated Na^+^ ions at the size exclusion filter, or 2) outward Na^+^ currents under reversed chemical gradients of Na^+^ and K^+^, at strongly positive membrane potentials to deter competing inward K^+^ flux. We tested these two predictions, first indeed observing the predicted robust inward currents in WT KCRs in the absence of extracellular K^+^, when V_m_ was < −80 mv. To test the second prediction, we performed electrophysiology after reversing the natural electrochemical gradients of Na^+^ and K^+^, that is, imposing high extracellular K^+^ concentration ([K^+^]_out_) and high intracellular Na^+^ concentrations ([Na^+^]_in_) (Figure 5E, left). WT *Hc*KCR1 maintained robust photocurrents under these new conditions (Figure S6C) but with E_rev_ shifted positively to 26.2 ± 3.3 mV under these conditions (Figure 5E, middle and right), revealing that Na^+^ indeed could efficiently move from the intracellular to the extracellular side and the proposed K^+^ selective filter could not prevent this from occurring. The same effect was observed in *Hc*KCR1 mutants with different ion selectivity or even *Hc*KCR2 WT and its mutants (Figure 5E, middle and right). Consistent with this interpretation, Attenuated Total Reflection Fourier-Transform InfraRed (ATR-FTIR) spectroscopy of WT *Hc*KCR1 showed that K^+^ does not stably bind to the selectivity filter both in dark and light conditions (Figures S3G and S3H). These results support the idea that the *Hc*KCR K^+^ selectivity filter does not tightly bind and specifically coordinate dehydrated K^+^ as observed in canonical K^+^ channels (Figure S1B) (Furutani et al., 2012), but instead favors flux of the smaller hydrated K^+^ ion at an aromatic size exclusion filter at the EV, and thus deters the entrance of Na^+^ ions (which would have to be chiefly from the extracellular side under physiological conditions, and which would encounter the presence of robust outward flux– and size filter occupancy– by smaller K^+^ ions).

We next performed molecular dynamics (MD) simulations of *Hc*KCR1 in the presence of K^+^. A series of 500-ns simulations provided three important findings (Figures 6A-C). First, K^+^ indeed does not stably bind at the EV but does occasionally bind spontaneously to a site near the IV, defined by the constriction formed at D116 and T120. Of note, these interactions are transient (Figure 6A, top), consistent with the ATR-FTIR result (Figure S3H). Second, these binding events are always accompanied by a loss of the salt bridge between R244 and D116 and reorientation of the R224 side chain towards the solvent. The K^+^ essentially replaces the guanidinium group of R224, making simultaneous binding unfavorable (Figure 6B). Third, when K^+^ binds to D116 and T120, some water molecules surrounding K^+^ in the solution are removed, rendering the K^+^ partially dehydrated (Figure 6C). These observations suggest that D116, and surrounding residues, present a favourable environment for partial dehydration of ions entering the cavity from the intracellular side. If this hypothesis is correct, the EV selectivity filter may also encounter partially dehydrated K^+^ ions arriving via the IV, which would also be good candidates for passing through an EV aromatic size exclusion filter since partially dehydrated K^+^ ions will have even smaller radii.

**Figure 6.**
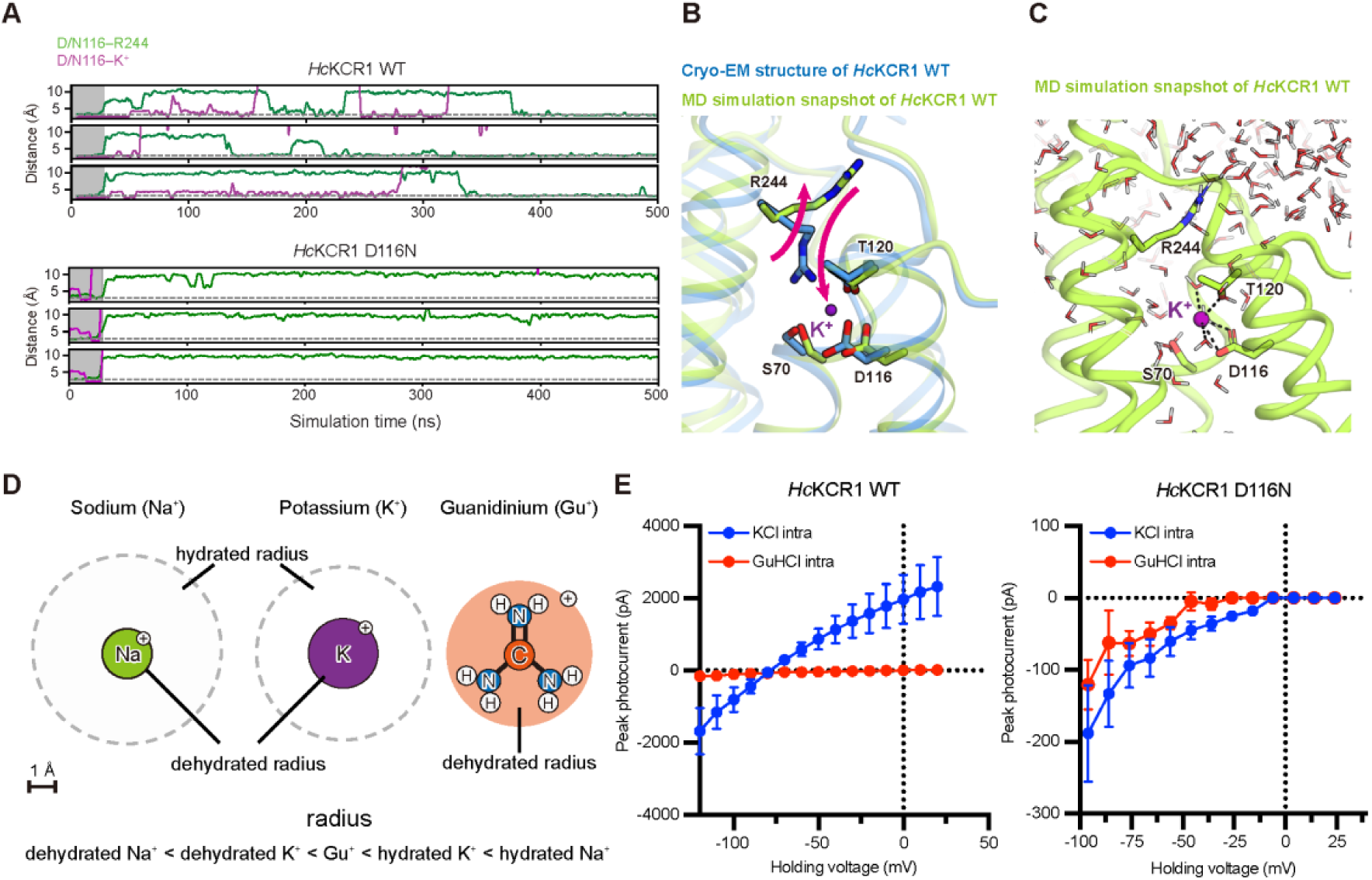
Computational and functional analyses of permeant ion dehydration. (A) Representative traces from molecular dynamics (MD) simulation of *Hc*KCR1 WT (top) and the D116N mutant (bottom); distances between D/N116 and R224 (green), and between D/N116 and K^+^ (magenta), are plotted for each monomer in a trimer. The gray shaded region at the beginning of the simulation marks the equilibration period during which the protein was restrained to the cryo-EM conformation. (B) Superposition of the *Hc*KCR1 cryo-EM structure and the MD simulation snapshot. The purple sphere indicates K^+^. Pink upward and downward arrows represent the flipping movements of R244 and the entry of K^+^ to the binding site, respectively. (C) MD simulation snapshot showing the transient binding of partially dehydrated K^+^. (D) The ionic and hydration radii of sodium (Na^+^), potassium (K^+^), and guanidinium (Gu^+^) ions. (E) Current-voltage (*I-V*) relationships of *Hc*KCR1 WT (left) and D116N mutant (right) in the presence and absence of GuHCl in the intracellular solution. Mean ± SEM (n = 3–8).

Computational and functional analyses of the D116N mutant were in good agreement with this hypothesis. MD simulation of the D116N mutant revealed that K^+^ does not bind to D116N and T120, unlike the case for WT *Hc*KCR1 (Figure 6A, bottom). Moreover, electrophysiology showed that currents were nearly abolished in this mutant, however, a small inward current of WT *Hc*KCR1 remained, reversing near 0 mV in this severe loss-of-function mutant and revealing that D116 is not absolutely required for ion conduction in general (Figure S5).

To further examine the importance of dehydration, we analysed the effect of guanidinium ions (Gu^+^) on the channel activity of *Hc*KCR1 (Figures 6D and 6E). Gu^+^ is a monovalent cation with radius larger than those of dehydrated K^+^ or Na^+^ but smaller than those of hydrated K^+^ or Na^+^ (Figure 6D); moreover, Gu^+^ is known to be one of the most weakly hydrated cations in solution (Mason et al., 2003). We found that addition of Gu^+^ to the intracellular solution completely inhibited channel activity of WT *Hc*KCR1 (Figure 6E, left). The lack of outward Gu^+^ current in itself indicated that either Gu^+^ acts as a pore blocker by interacting with a specific binding site in the pore, or that Gu^+^ is simply too large an ion for *Hc*KCR1 to transport (in which case even larger cations– such as fully hydrated K^+^ or Na^+^– would also be too large for *Hc*KCR1 to transport, suggesting that partial dehydration may be important for ion transport). We next found that Gu^+^ indeed blocked the transport of K^+^ and Na^+^ (Figure 6E, left). Considering the structural similarity between Gu^+^ and the side chain of the arginine residue, it is possible that D116 serves as a Gu^+^ binding site as R244 interacts with D116, and that Gu^+^ binding to this site prevents ion flux. This idea was supported by further electrophysiology of the D116N mutant showing that Gu^+^ does not significantly inhibit the inward current remaining in the mutant (Figure 6E, right), presumably because it is no longer able to bind to D116 just as is the case with the R244 guanidinium moiety.

### A Proposed Mechanism for the High K^+^ Permeability Ratios of KCRs

Our structural, electrophysiological, spectroscopic, and computational data collectively provide insights into the mechanism for the high K^+^ permability ratios of KCRs. Under physiological conditions, the concentrations of most simple cations, including Na^+^, Ca^2+^, and Mg^2+^, are higher on the extracellular side, while the concentration of K^+^ is higher on the intracellular side. When the KCR is opened by light, fully-hydrated Na^+^ approaching from the extracellular side encounters a barrier at the size selectivity filter formed by W102, F/Y221, Y222, and H225 (hydrated Ca^2+^ and Mg^2+^, even larger than hydrated Na^+^, would be blocked by the same mechanism); moreover hydrated Na^+^ is outcompeted by 1) smaller hydrated K^+^ ions from the extracellular side, if present, and 2) even smaller partially dehydrated K^+^ ions from the intracellular side (Figure 7A) where R244 is mobile; when it flips out intracellular K^+^ can bind to D116, stripping away some water molecules in the hydration shell.

**Figure 7.**
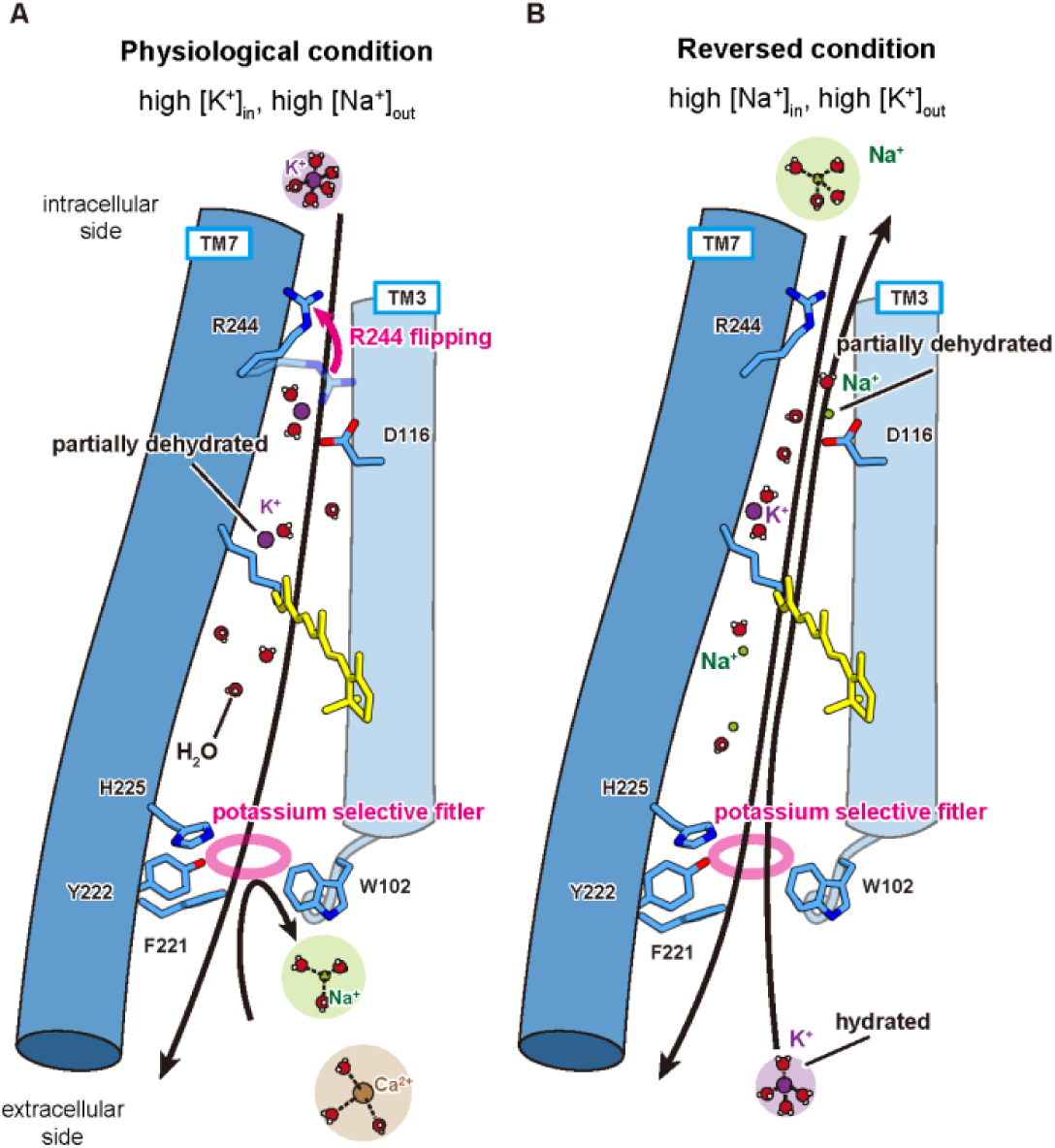
Proposed model for high K^+^ permeability ratios in KCRs. (A) When the concentrations of extracellular Na^+^ and intracellular K^+^ are high (physiological/normal condition), permeation of large, hydrated cations such as Na^+^ and Ca^2+^ is blocked at the size filter formed by W102, F/Y221, Y222, and H225. In contrast, K^+^ can enter the pore (under physiological ion balance conditions chiefly from the intracellular side, when the interaction between D116 and R244 is broken); K^+^ can become partially dehydrated, permeate through the ion-conducting pathway, and pass the size filter for release to the extracellular space. (B) When the concentrations of intracellular Na^+^ and extracellular K^+^ are high (reversed condition), Na^+^ can move outward, just as K^+^ moves outward under physiological conditions. Inward K^+^ currents are possible in this condition through the aromatic size-exclusion filter at the EV (the size of hydrated K^+^ is smaller than that of Na^+^ or Ca^2+^). TMs 1, 2, 4-6 are removed for clarity. Black and pink arrows indicate the cation flow and the conformational change of R244, respectively. K^+^, Na^+^, Ca^2+^, and oxygen and hydrogen atoms of water molecules are shown as spheres colored in purple, green, brown, red and small white, respectively. Magenta circles represent the K^+^ selective size filters.

A similar route would also be available for intracellular Na^+^, including partial dehydration at D116, but would rarely be taken under physiological conditions due to very low [Na^+^]_i_. However, Na^+^ currents can be observed, especially if K^+^ flux is prevented by manipulating electrochemical gradients. By completely removing extracellular K^+^ (Fig. 5E), we enabled inward Na^+^ currents at V_m_ < −80 mV (revealing a phenomenon which can be observed in canonical K^+^ channels as well, such as Kv2.1– namely K^+^ block of K^+^ channels, specifically the block of non-K^+^ flux through K^+^ channels by K^+^ ions that compete more effectively at the selectivity filter) (e.g. Kiss et al., 1998). Consistent with this interpretation, when we artificially reversed the concentrations of intracellular K^+^ and extracellular Na^+^, the direction of ion flow was also reversed (Figure 7B). Notably, presumably-hydrated K^+^ can still efficiently flow from the extracellular to intracellular side in this condition (Figures 6B and S6C), suggesting that the size boundary for ionic species that can pass through the selectivity filter is above that of hydrated K^+^.

In conclusion, KCRs adopt a unique mechanism to specifically favor K^+^ flux, in a manner unlike canonical K^+^ channels. The KCR channels employ an aromatic size exclusion filter for hydrated or partially-hydrated ions at the extracellular side, rather than specifically coordinating dehydrated ions. Species with small, hydrated radii are favored for transport, a process dominated by outward K^+^ flux under physiological conditions due to the strong electrochemical gradient in that direction.

## DISCUSSION

Because of high K^+^ permeability ratios as well as robust conductance and light sensitivity (Govorunova et al., 2022), KCRs have attracted much attention as potential tool for optogenetic neuronal silencing. However, the mechanisms underlying these remarkable properties had not been understood. Here, we have revealed the structural basis for these key properties, along with insights into the evolution of K^+^ permeability ratios in microbial opsins.

Our *Hc*KCR1 and *Hc*KCR2 structures represent the second and third high-resolution structures of proteins in the PLCR family, allowing for the first time a structural comparison among PLCRs. We found that the most of the unique features observed in the structure of ChRmine, the first structure from the PLCR family (Kishi et al., 2022), are conserved in both *Hc*KCR1 and *Hc*KCR2 – trimeric assembly, short TM3, deformed Schiff base, and large cavities within the monomer (Figures 1-4) – suggesting that these features may be typical for the PLCR family.

However, detailed comparisons reveal several key differences. For example, the primary counterion and water distribution in the Schiff base region are different between *Hc*KCR1 and ChRmine, which may be related to the different compositions of the retinal isomers after light absorption; in ChRmine, as in other microbial rhodopsins, all-*trans* retinal is isomerized mostly to 13-*cis* upon light illumination (Kishi et al., 2022), while in *Hc*KCR1, the percentage of increased 9-, 11-, and 13-*cis* retinal are about the same (Figures S3A and S3B). The 13-*cis* retinal-bound *Hc*KCR1 likely corresponds to the open state, but further studies will be needed to confirm this and to identify the function of the 9- and 11-*cis* retinal-bound photoproducts.

The retinal isomer is different even in the dark condition between *Hc*KCR1 and 2. Unlike most ChRs including *Hc*KCR1, *Hc*KCR2 has a twisted 6-s-*cis* form of the retinal isomer (Figure 3E). Only two residues (T136/G140 in *Hc*KCR1 and A136/A140 in *Hc*KCR2) are involved in the conversion from 6-s-*trans* to 6-s-*cis*, and considering findings from a previous study of C1C2 (Kato et al., 2015a), G-to-A replacement at position 140 of *Hc*KCR1 might be sufficient for the conversion, and the residue at position 136 may be important to finely tune the amount of twist. Interestingly, while the glycine at position 140 of *Hc*KCR1 is highly conserved among microbial rhodopsins (Figure S1A), all blue-shifted ChRs, including *Ps*ChR, *Ts*ChR, *Kn*ChR, WiChR, and *B1*ChR2, have alanine at this position (Govorunova et al., 2013; Klapoetke et al., 2014; Tashiro et al., 2021; Vierock et al., 2022). Therefore, we believe that this “non-G rule” is generalizable and may prove useful in the search for microbial rhodopsins with 6-s-*cis* retinal and blue-shifted absorption spectra.

The most notable difference between *Hc*KCR and ChRmine is observed in the interaction network at the ECS (Figure 4). Because of the displacements of ECL1 and TM7 (Figure 1F), four aromatic residues on the helix and loop (W102, F/Y221, Y222, and H225) can assemble and form the K^+^ selectivity filter in *Hc*KCRs (Figure 4A). However, the interaction network is not identical even between *Hc*KCR1 and 2 (Figures 4C and S2W). The replacements of several residues including D/N18, L/Q211, Q/R218, and F/Y221, make the environment around the ECS slightly different (Figure S1A); this difference could contribute to the different K^+^ permeability ratios between *Hc*KCR1 and 2 (*P*_K_/*P*_Na_ values of *Hc*KCR1 and 2 are 25.7 ± 2.7 and 17.0 ± 1.5, respectively). Notably, a recently discovered KCR with higher K^+^ permeability ratio, WiChR, has different amino acids at positions 18, 210, 211, 218, 221, and 222 (*Hc*KCR1 numbering), suggesting that the interaction network at the ECS is also different in WiChR and obtaining a high-resolution structural information of WiChR would be of enormous value to understand the difference in K^+^ selectivity within KCRs. More detailed comparison and further understanding of the working mechanism of the filter might lead to the development of novel KCRs with improved K^+^ selectivity; indeed we can already successfully create *Hc*KCR mutants with *P*_K_/*P*_Na_ value of >50, such as H225F, using information from our *Hc*KCR structures (Figure 5B).

The H225F mutant shows not only a high K^+^ permeability ratio (*P*_K_/*P*_Na_ = 53.9 ± 4.6) but also comparable photocurrent amplitude to WT (Figures 5B, S5, and S6A), suggesting usefulness as a novel tool for optogenetics. In addition to H225F, we identified two more mutations that enhance the properties of *Hc*KCRs: C110T and Y222A (Figures 3C, 3D, 5B, S5, and S6A). It was a surprise that the C110T mutation increases the τ_off_ of *Hc*KCRs by more than 1500-fold, because previous attempts to introduce the same mutation to other PLCRs failed. The transferability of this mutation may be higher than previously expected (Sineshchekov et al., 2020), and so more SFO mutants of PLCRs now can be expected in the future. On the other hand, the Y222A mutation does not significantly affect τ_off_ but E_rev_ is depolarized by ∼70 mV, converting the function of *Hc*KCRs from inhibitory to excitatory channels under physiological conditions. The photocurrent amplitude of this mutant is comparable to or even higher than that of WT *Hc*KCR (Figures S5 and S6A); thus, this novel KCR could be useful as a potent excitatory optogenetics tool.

As far as we know, all canonical K^+^ channels discovered and structurally resolved (until this work) have shared the same basic tetrameric assembly and K^+^ selectivity filter (González et al., 2012) (Figure S1B). Of note, a lysosomal ion channel TMEM175, which was initially reported as a K^+^-selective channel (Cang et al., 2015; Lee et al., 2017; Oh et al., 2020) but later shown to be more H^+^ selective (Hu et al., 2022), lacks the K^+^ selectivity filter but shares the tetrameric assembly property. Interestingly, microbial rhodopsins are known to form diverse oligomeric assemblies, but these are all dimers, trimers, pentamers, or hexamers, and not a single rhodopsin forming a tetramer has been reported (Nagata and Inoue, 2021). This observation suggests that tetrameric assembly may be unstable for microbial rhodopsins, which achieve pore selectivity within each 7TM monomer. A previous study suggests that the orientation of the conserved arginine residue on TM3/ECL1 (R82 in *Hs*BR and R112 in ChRmine) may be important to define channel-vs. pump-type rhodopsins, and indeed replacement of this arginine to glutamine significantly affects function and ion selectivity of outward Na^+^ pump rhodopsin KR2 (Figure S4C) (Kishi et al., 2022; Vogt et al., 2019). In KCRs, this arginine is replaced by tryptophan, and we suggest that this replacement would be an important step in evolving high K^+^ permeability ratios.

To fully understand the KCR photocycle, further studies, including structural determination of intermediate states in *Hc*KCRs, will be needed. However, our current analysis has already revealed key mechanisms underlying high K^+^ permeability ratios in KCRs and achieved molecular engineering of several new KCR variants with improved functionality: C110T, Y222A, and H225F. These findings provide insight into the evolution of K^+^ channels and rhodopsins, and enable the discovery and development of new opsins with distinct ion selectivity. These opsins, together with the variants we have developed in this study, may further diversify and improve optogenetic technologies, opening up new avenues for basic life science and biomedical research.

## ACKNOWLEDGEMENTS

We thank H. Yasumoto, K. Hasegawa (Univ. of Tokyo), and C. Delacruz (Stanford Univ.) for administrative support, and T. Kawamura (Univ. of Tokyo) for technical support. This work was supported by the MRC, as part of UKRI (MC_UP_A025_1012 to K.Y.), The Nakatani Foundation (H.E.K.), The Mitsubishi Foundation (H.E.K.), The Kazato Research Foundation (H.E.K.), the UTEC-UTokyo FSI Research Grant Program (H.E.K.), AMED (JP21wm0525018 to H.E.K.), JSPS KAKENHI (22H04742 to M.F., JP20K21383/JP21H01875 to K.I., and 21H05142/22H00400/22K19265 to H.E.K.), JST SPRING (JPMJSP2108 to S.T.), JST PRESTO (JPMJPR1888 to T.N.), JST FOREST (JPMJFR204S to H.E.K.), JST CREST (JPMJCR21P3 to H.E.K.), the NSF NeuroNex program (K.D.), the NOMIS Foundation (K.D.), the Else Kröner Fresenius Foundation (K.D.), the Gatsby Foundation (K.D.), and a grant for ChR structure determination from the NIMH (R01MH075957 to K.D.).

## AUTHOR CONTRIBUTIONS

S.Tajima expressed and purified the proteins and prepared cryo-EM grids with the help of M.F. and K.E.K. S.Tajima and M.F. obtained cryo-EM images. S.Tajima and T.E.M. processed cryo-EM data to generate 3D maps. S.Tajima, M.F., K.E.K., Y.K., and H.E.K. built the models and refined the structures. Y.S.K., E.F.X.B., and P.Y.W. performed the electrophysiology with support of C.R. J.M.P. performed and analyzed the molecular dynamics simulations under the supervision of R.O.D. C.R. and H.I. carried out the molecular cloning and mutagenesis. S.Tajima measured UV-Vis absorption spectra. M.K. and K.I. performed flash photolysis. T.N. performed HPLC analysis to determine the retinal isomer. S.Takaramoto performed the laser patch clamp experiment. M.S. and K.K. performed the ATR-FTIR experiment under the supervision of H.K. M.I. provided input on the manuscript. S.Tajima, Y.S.K., K.D., and H.E.K. wrote the manuscript with input from all the authors. K.D. and H.E.K. supervised all aspects of the research.

---

**Figure S1.**
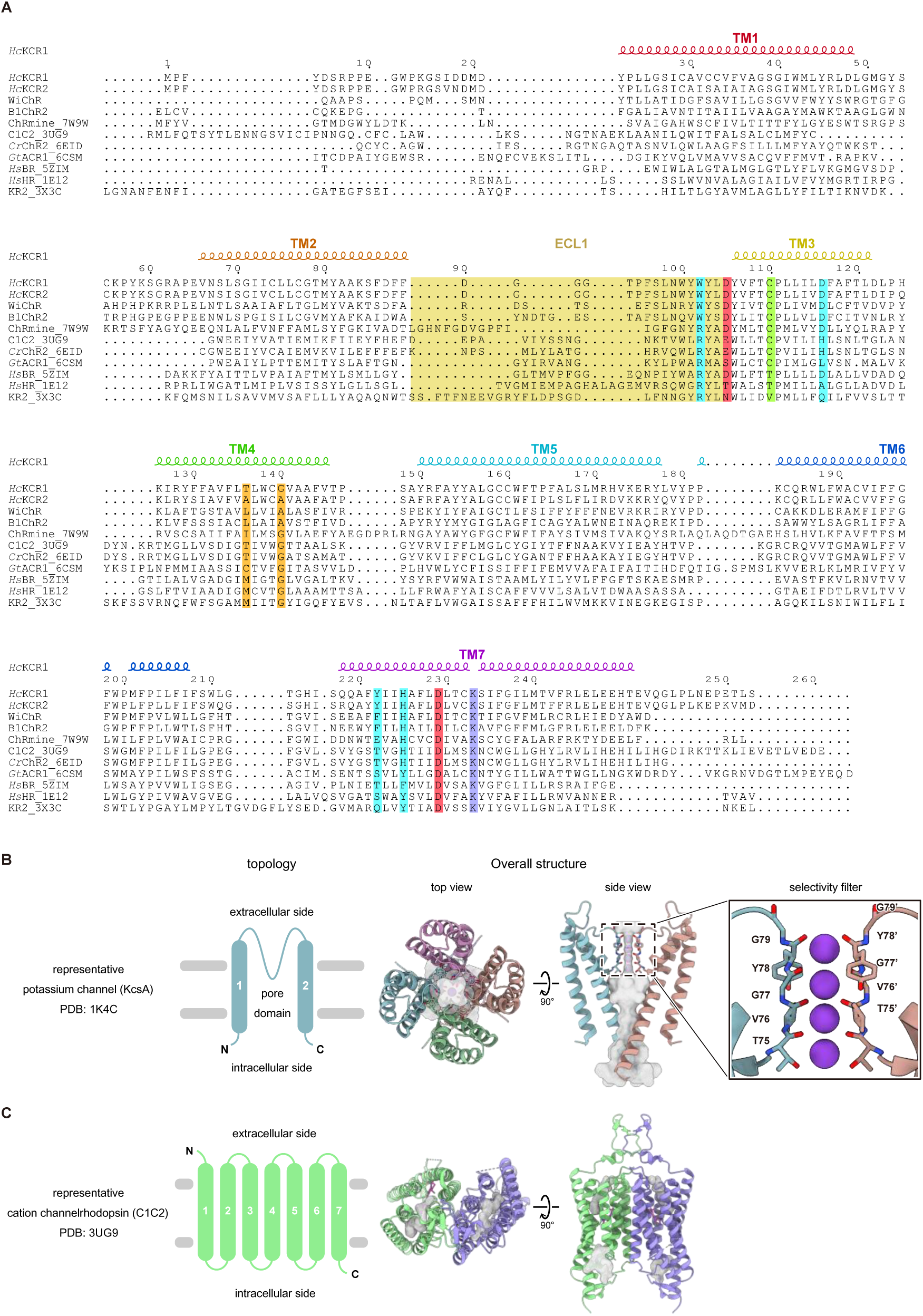
Comparisons of sequence, topology, and assembly, related to Figure 1. (A) Structure-based amino acid sequence alignment of microbial rhodopsins. The sequences are *Hc*KCR1 (GenBank: MZ826862), *Hc*KCR2 (GenBank: MZ826861), WiChR (Vierock et al., 2022), ChRmine (PDB: 7W9W) (Kishi et al., 2022), C1C2 (PDB: 3UG9)(Kato et al., 2012), *Cr*ChR2 (PDB: 6EID) (Volkov et al., 2017), *Gt*ACR1 (PDB: 6CSM) (Kim et al., 2018), *Hs*BR (PDB: 5ZIM) (Hasegawa et al., 2018), *Hs*HR (PDB: 1E12) (Kolbe et al., 2000), and KR2 (PDB: 3X3C) (Kato et al., 2015b). The sequence alignment was created using PROMALS3D (Pei et al., 2008) and ESPript 3 (Robert and Gouet, 2014) servers. Secondary structure elements for *Hc*KCR1 are shown as coils. The lysine forming the Schiff base with retinal is colored in purple. The cysteine for the step-function variant is colored in green. The counterion candidates are colored in red. The ECL1 regions are highlighted in pale yellow. The residues forming the pocket for the β-ionone ring are colored in orange. The residues forming the dehydration gate and K^+^ selectivity filter are colored in cyan. (B) Architecture of the representative prokaryotic K^+^ channel, KcsA (PDB: 1K4C). Transmembrane topology (left). Each subunit contains two TMs with a short loop containing the K^+^ selectivity filter. The tetrameric assembly viewed from the extracellular side and viewed parallel to the membrane (middle), colored by protomer (blue, green, red, and orange). The ion-conducting cavity is colored in semitransparent grey. K^+^ ions and the TVGYG motif are depicted by ball and stick models, respectively. Magnified view of the selectivity filter (right). Only two subunits are shown for clarity. (C) Architectures of the representative channelrhodopsin, C1C2 (PDB: 3UG9). Transmembrane topology (left). Each subunit contains seven TMs without any TVGYG or related motif. The dimeric assembly viewed from the extracellular side and viewed parallel to the membrane (right), colored by protomer (green and purple). The ion-conducting cavity is colored in grey.

**Figure S2.**
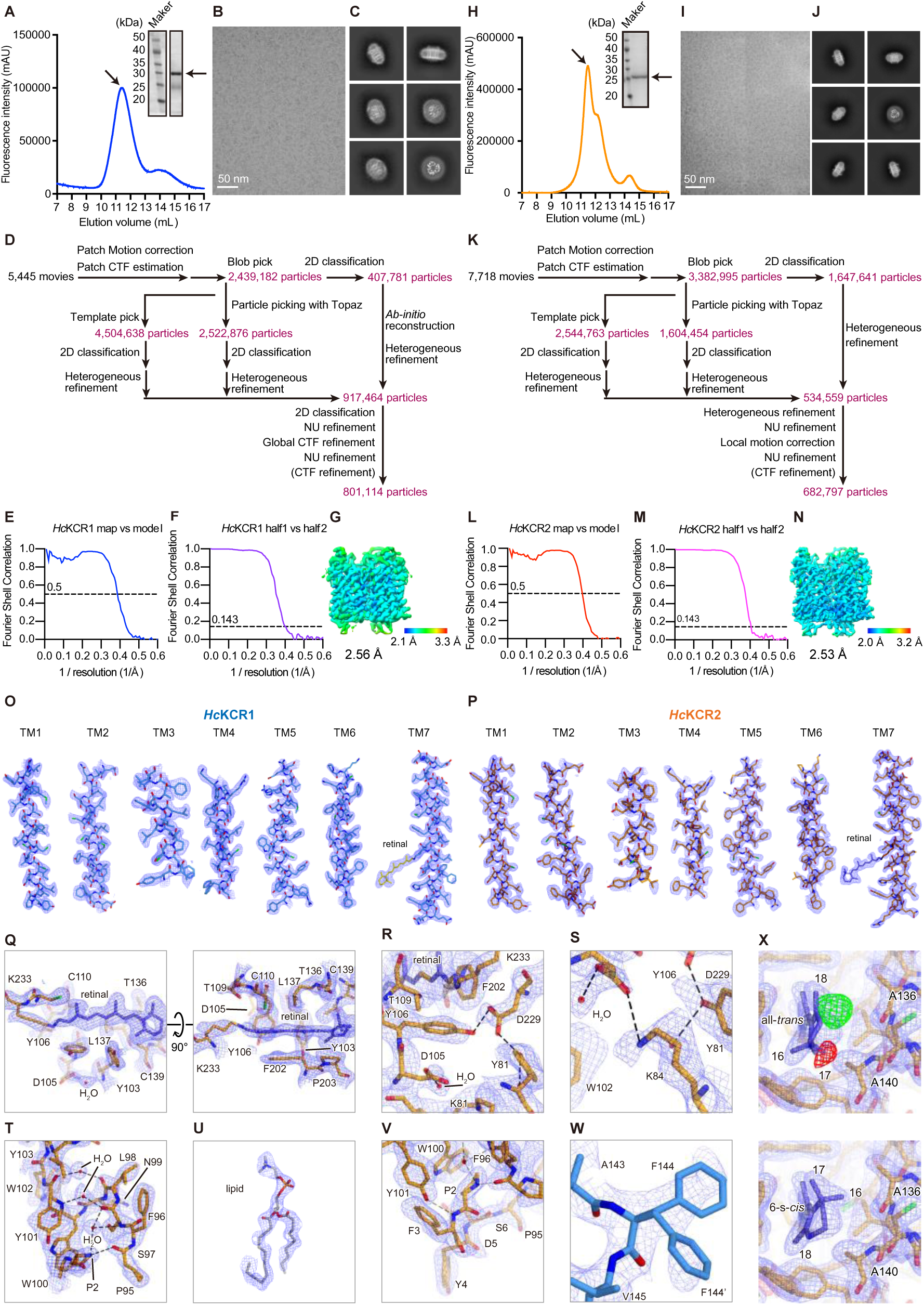
Cryo-EM analysis of the *Hc*KCR1 and *Hc*KCR2, related to Figures 1-3. (A–G) Panels corresponding to *Hc*KCR1 reconstituted into MSP1D1E3. (A) Representative SEC trace with SDS-PAGE as inset. The fluorescence signals from tryptophan residues were monitored by the fluorescence detector (excitation: 280 nm, emission: 350 nm). Black arrows represent the *Hc*KCR1 protein. (B) Representative cryo-EM micrograph. (C) 2D-class averages. (D) Data processing workflow of the *Hc*KCR1 reconstituted into MSP1D1E3. (E) Fourier shell correlation (FSC) between the two independently refined half-maps. (F) FSC between the model and the map calculated for the model refined against the full reconstruction. (G) Final cryo-EM map colored by local resolution. (H–N) Panels corresponding to *Hc*KCR2 reconstituted into MSP1D1E3. (H) Representative SEC trace with SDS-PAGE as inset. Black arrows represent the *Hc*KCR2 protein. (I) Representative cryo-EM micrograph. (J) 2D-class averages. (K) Data processing workflow of the *Hc*KCR2 reconstituted into MSP1D1E3. (L) FSC between the two independently refined half-maps. (M) FSC between the model and the map calculated for the model refined against the full reconstruction. (N) Final cryo-EM map colored by local resolution. (O–W) Representative cryo-EM densities of *Hc*KCR1and *Hc*KCR2. FSC-weighted sharpened maps were calculated by cryoSPARC v3.2.0 for *Hc*KCR1 and cryoSPARC v3.3.2 for *Hc*KCR2, respectively. Transmembrane helices for *Hc*KCR1 (O) and *Hc*KCR2 (P). Retinal binding pocket (Q), the Schiff base region (R), K84 (S), ECL1 (T), lipid molecule (U), the N-terminal region (V) of *Hc*KCR2. (W) Two different rotamers observed in F144 of *Hc*KCR1. Stick models are colored in blue for *Hc*KCR1 and orange for *Hc*KCR2, respectively. (X) Cryo-EM densities around the β-ionone ring of *Hc*KCR2. Blue and green/red maps are FSC-weighted sharpened map calculated by cryoSPARC v3.3.2, and *F*_o_-*F*_c_ maps calculated by the program Servalcat, respectively. All-*trans* (top) and 6-s-*cis* (bottom) retinal are modeled against the FSC-weighted sharpened map. Positive (green) and negative (red) *<u>F</u>*_o_-*F*_c_ difference density pairing (±5.2 σ, where σ is the standard deviation within the mask) is observed between C_18_ and A136 (top), suggesting rotation of the β-ionone ring.

**Figure S3.**
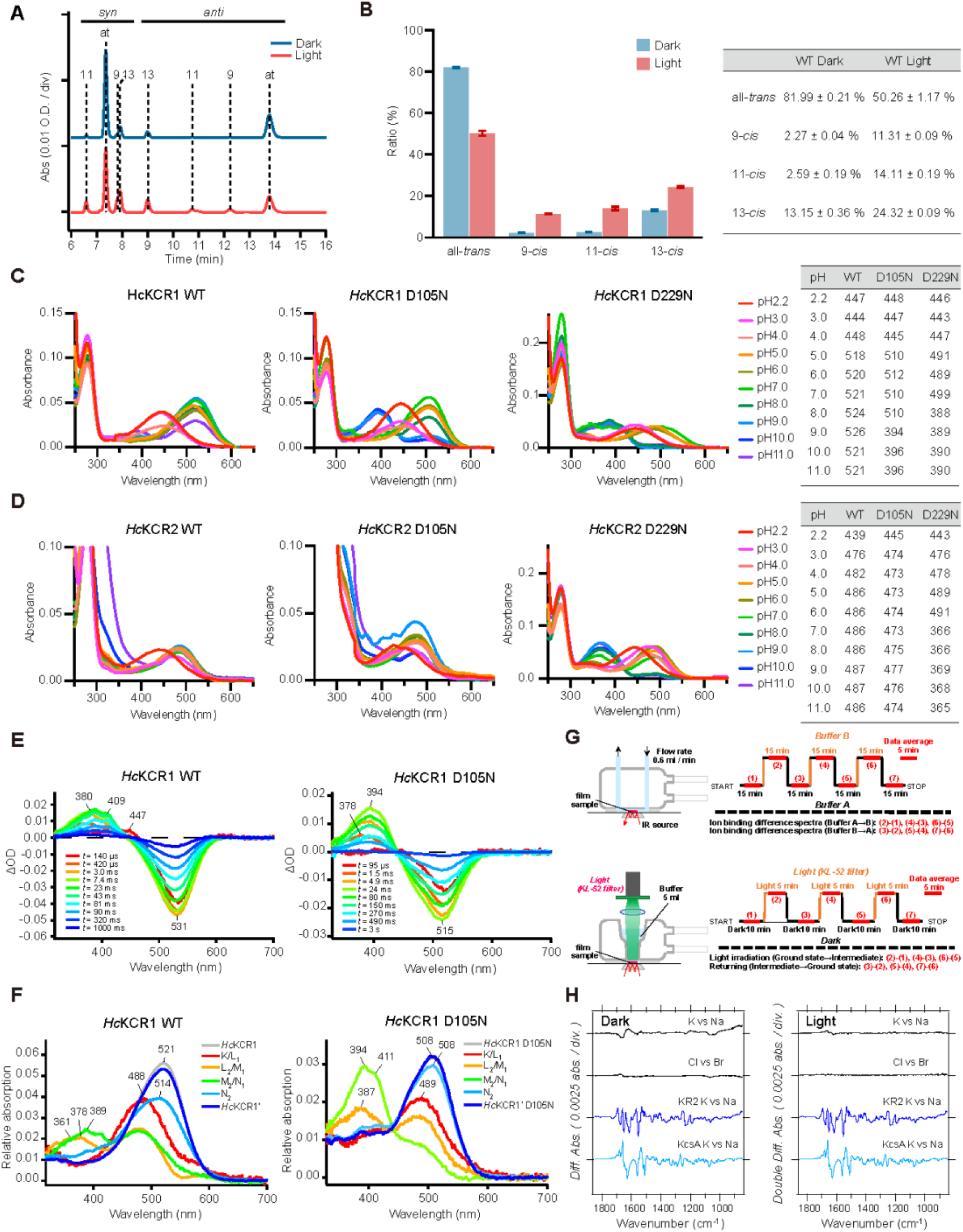
Spectroscopic and HPLC characterization of *Hc*KCR1 and *Hc*KCR2, related to Figures 1, 2, and 5. (A and B) HPLC analysis of the chromophore configuration of *Hc*KCR1 WT. (A) Representative HPLC profiles of the chromophore of *Hc*KCR1 under dark (top) and light conditions (bottom). Abbreviations ‘‘at’’, ‘‘9’’, ‘‘11’’, and ‘‘13’’ indicate the peaks of all-*trans*, 9-*cis*, 11-*cis*, and 13-*cis* retinal oximes, respectively. (B) Calculated composition of retinal isomers in *Hc*KCR1 under dark and light conditions. Data are presented as mean ± SEM (n = 3). Values are listed in the table. Green light (510 ± 5 nm) was used for illumination. Light adaptation was achieved by illumination for 1 min followed by incubation in the dark for 2 min. (C and D) pH-titrated absorption spectra of *Hc*KCR1 and *Hc*KCR2. (C) The absorption spectra of *Hc*KCR1 WT (left), D105N (middle), and D229N (right) from pH 2.2 to 11.0. (D) The absorption spectra of *Hc*KCR2 WT (left), D105N (middle), and D229N (right) from pH 2.2 to 11.0. The λ_max_ value at each pH is listed in the table. (E) Transient absorption spectra of *Hc*KCR1 WT (left) and D105N (right). (F) The absorption spectra of the initial state (gray), K/L_1_ (red), L_2_/M_1_ (orange), M_2_/N_1_ (green), N_2_ (light blue), and *Hc*KCR1ʹ of *Hc*KCR1 WT (left), and those of the initial state (gray), K/L_1_ (red), L_2_/Mʹ (orange), Mʹʹ /N_1_ (green), N_2_ (light blue), and *Hc*KCR1ʹ of D105N (right). The spectra are calculated from the decay-associated spectra of transient absorption changes shown in Figures 2D and S3E. (G) Experimental procedure of ATR-FTIR spectroscopy either ion perfusion (top) or light illumination (bottom) systems. (H) ATR-FTIR difference spectra upon exchange of NaCl/KCl (top), NaCl/NaBr (middle) for *Hc*KCR1 in dark (left) and light (right) conditions. The spectra for KR2 are shown in the bottom as the reference. Unlike the case with KR2 and KcsA, the flat spectra of *Hc*KCR1 indicate that K^+^ does not stably bind to *Hc*KCR1 in either dark or light conditions.

**Figure S4.**
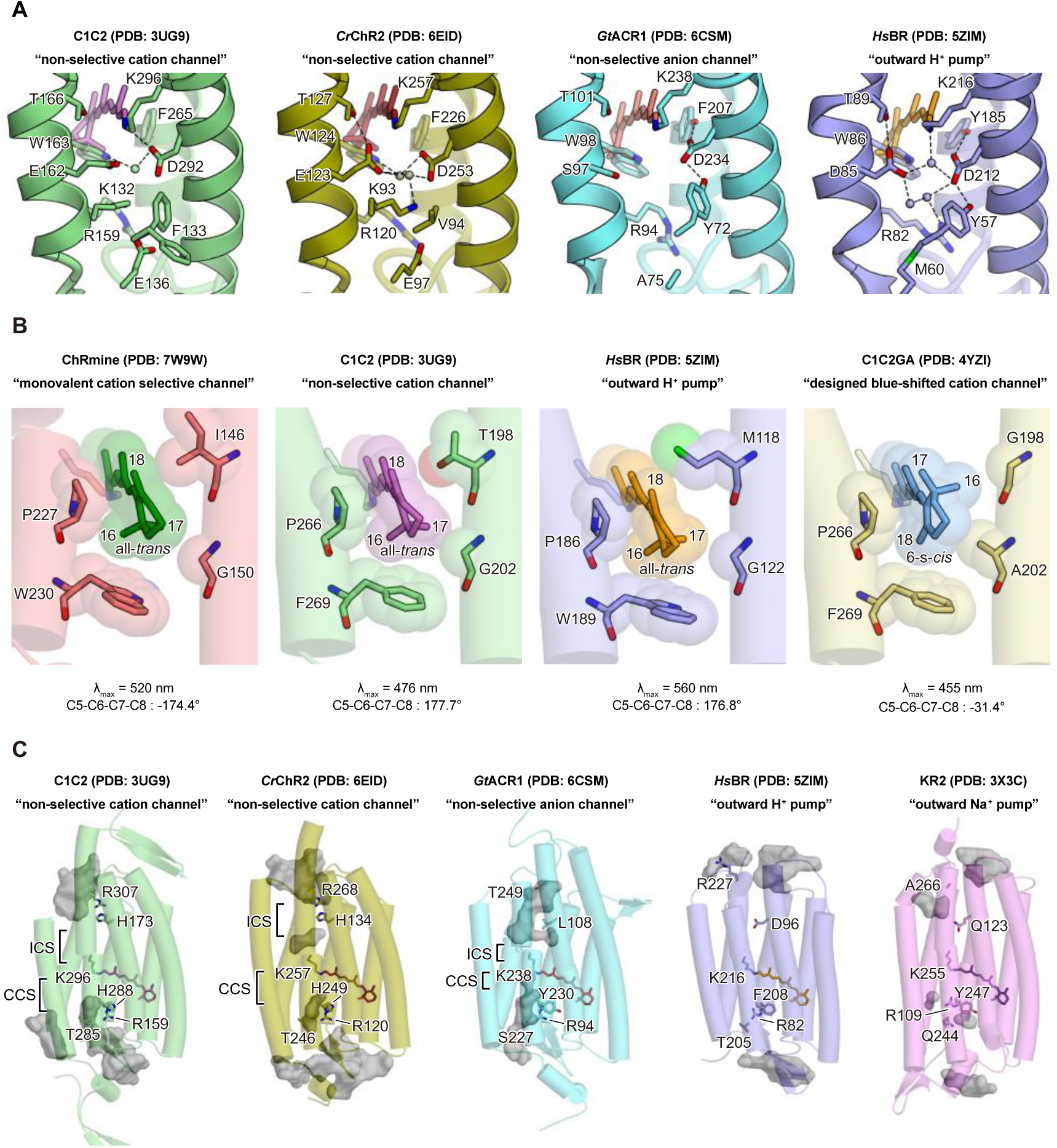
Structural comparison of representative microbial rhodopsins, related to Figures 2-4. (A) The Schiff base regions of C1C2 (PDB: 3UG9), *Cr*ChR2 (PDB: 6EID), *Gt*ACR1 (PDB: 6CSM), and *Hs*BR (PDB: 5ZIM). Spheres represent water molecules. The black dashed lines indicate H-bonds. (B) Retinal β-ionone ring in ChRmine (PDB: 7W9W), C1C2 (PDB: 3UG9), *Hs*BR (PDB: 5ZIM), and C1C2GA mutant (PDB: 4YZI). (C) Ion-conducting pathways in C1C2 (PDB: 3UG9), *Cr*ChR2 (PDB: 6EID), *Gt*ACR1 (PDB: 6CSM), *Hs*BR (PDB: 5ZIM), and KR2 (PDB: 3X3C). Key residues for K^+^ selectivity in *Hc*KCRs are shown as stick models. Intra- and extracellular cavities are calculated with the program HOLLOW.

**Figure S5.**
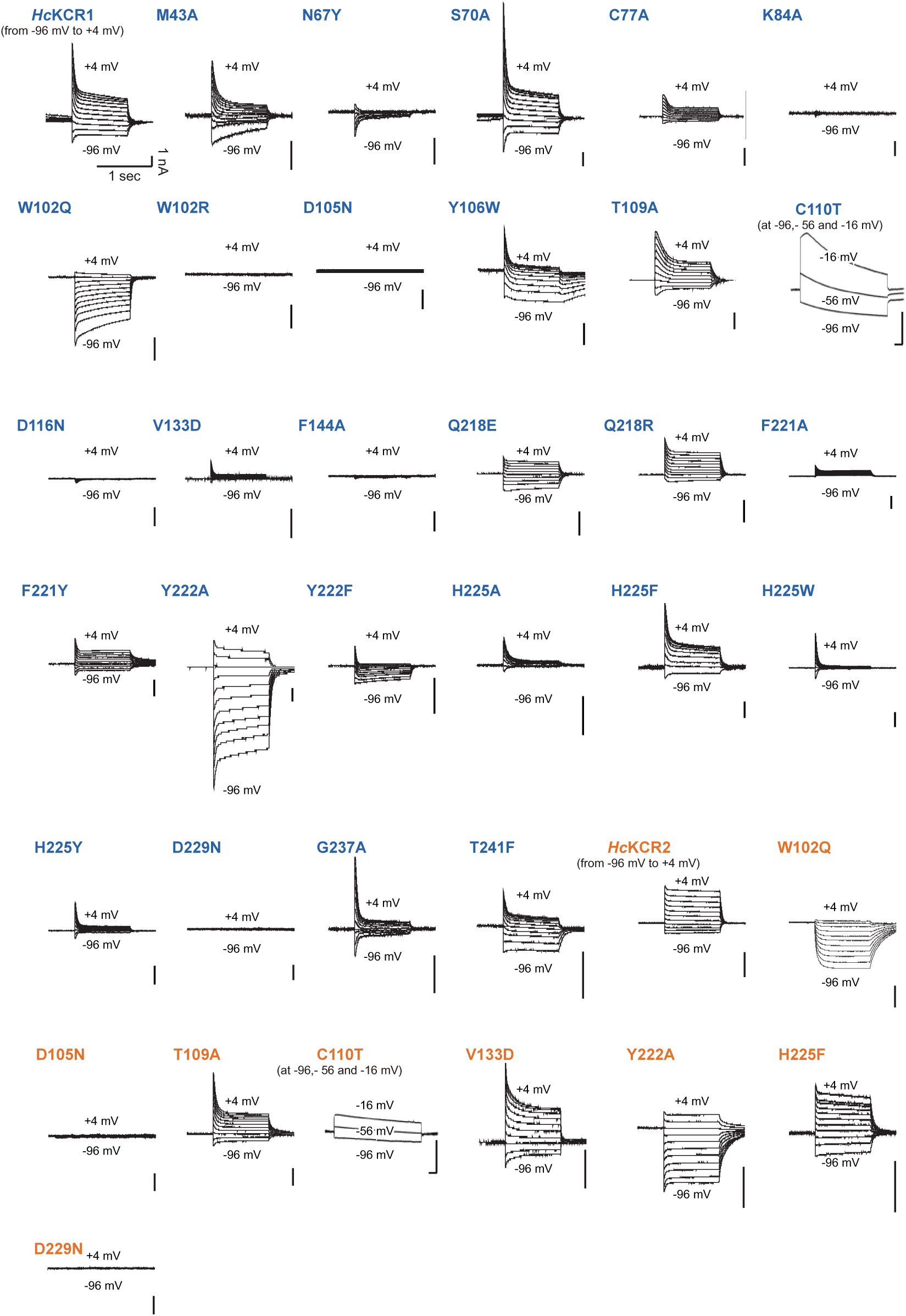
Representative traces for current–voltage measurements., related to Figures 2, 3, and 5. Voltage-clamp traces of *Hc*KCR1 WT and 27 mutants (colored in blue) and *Hc*KCR2 WT and 8 mutants (colored in orange), collected from −96 mV to +4 mV in steps of 10 mV (for C110T mutants, traces are collected from −96, −56, and −16 mV). HEK293 cells were recorded while stimulated by 1 s of 0.7 mW mm^−2^ irradiance at 560 nm for *Hc*KCR1 and 470 nm for *Hc*KCR2.

**Figure S6.**
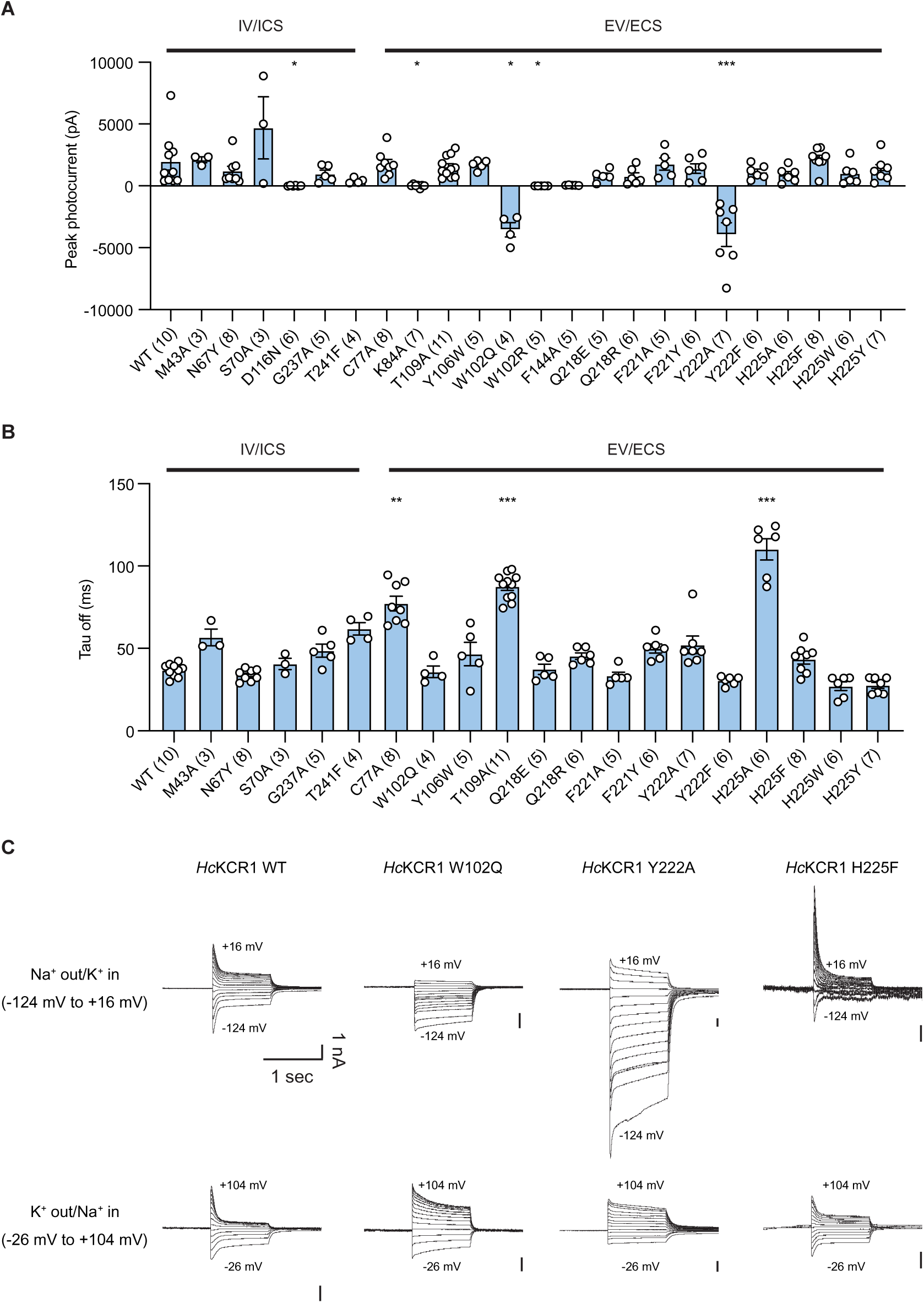
Electrophysiology, related to Figure 5. Summary of the peak photocurrent (A) and τ_off_ of channel closing (B). Mutants are categorized by location: intracellular vestibule or internal constriction site (IV/ICS) vs. extracellular vestibule or extracellular constriction site (EV/ECS). Sample size (number of cells) indicated in parentheses. Data are mean ± S.E.M (n = 3–11); one-way ANOVA followed by Dunnett’s test. * p < 0.05, ** p < 0.01, *** p < 0.001, and **** p < 0.0001. (C) Voltage-clamp traces of *Hc*KCR1 WT and 3 mutants in physiological (top) and reversed (bottom) conditions. Traces are collected from −124 mV to +16 mV in steps of 10 mV for the physiological condition and from −26 mV to +104 mV in steps of 10 mV for the reversed condition. HEK293 cells were recorded while stimulated with 1 s of 0.7 mW mm^−2^ irradiance at 560 nm.

## STAR★METHODS

### KEY RESOURCES TABLE

| REAGENT or RESOURCE | SOURCE | IDENTIFIER |
| --- | --- | --- |
| Bacterial and virus strains |  |  |
| One shot Stbl3 e-coli | Thermo Fisher Scientific | Cat# C737303 |
| *E. coli* BL21 (DE3) | Thermo Fisher Scientific | Cat# EC0114 |
| *E. coli* Mach1 | Thermo Fisher Scientific | Cat# C862003 |
| *E. coli* DH10Bac | Thermo Fisher Scientific | Cat# 10361012 |
| Chemicals, peptides, and recombinant proteins |  |  |
| Lipofectamine 2000 | Thermo Fisher Scientific | Cat# 11668027 |
| FuGENE | Promega | Cat# E2311 |
| Benzamidine | Nacalai tesque | Cat# M9G4533 |
| Leupeptin | Peptide Institute | Cat# 4041 |
| *n*-Dodecyl-β-D-maltoside (DDM) | EMD Millipore | Cat# 324355 |
| Ni-NTA Superflow | QIAGEN | Cat# 30430 |
| Superdex 200 Increase 10/300 GL | Cytiva | Cat# 28990944 |
| All-*trans*-retinal (ATR) | Sigma-Aldrich | Cat# R2500 |
| StockOptions pH buffer kit | Hampton Research | Cat# HR2-241 |
| ESF 921 Insect Cell Culture Medium, Protein Free | Expression systems | Cat# 96-001-01 |
| Deposited data |  |  |
| HcKCR1 – atomic model | From this paper | PDB: XXXX |
| HcKCR2 – atomic model | From this paper | PDB: XXXX |
| HcKCR1 – Cryo-EM map | From this paper | EMDB-XXXXX |
| HcKCR2 – Cryo-EM map | From this paper | EMDB-XXXXX |
| HcKCR1 – Cryo-EM movie frames | From this paper | EMPIAR-XXXXX |
| HcKCR2 – Cryo-EM movie frames | From this paper | EMPIAR-XXXXX |
| Experimental models: Cell lines |  |  |
| *Spodoptera frugiperda* Sf9 cells | Expression systems | Cat# 94-001F |
| HEK293T cells | ATCC | Cat# CRL-3216 |
| Recombinant DNA |  |  |
| pFastBac-HcKCR1 (WT)-EGFP-His10 | This paper | N/A |
| pFastBac-HcKCR1 (D105N)-EGFP-His10 | This paper | N/A |
| pFastBac-HcKCR1 (D229N)-EGFP-His10 | This paper | N/A |
| pFastBac-HcKCR1 (T136A/G140A)-EGFP-His10 | This paper | N/A |
| pFastBac-HcKCR2 (WT)-EGFP-His8 | This paper | N/A |
| pFastBac-HcKCR2 (D105N)-EGFP-His8 | This paper | N/A |
| pFastBac-HcKCR2 (D229N)-EGFP-His8 | This paper | N/A |
| pFastBac-HcKCR2 (A136T/A140G)-EGFP-His8 | This paper | N/A |
| pET-43a(+)-MSP1D1E3 | This paper | N/A |
| pcDNA 3.1-HcKCR1 (WT)-eYFP | This paper | N/A |
| pcDNA 3.1-HcKCR2 (WT)-eYFP | This paper | N/A |
| Software and algorithms |  |  |
| Serial EM software | (Mastronarde, 2003) | https://bio3d.colorado.edu/SerialEM/ |
| MotionCor2 | (Zheng et al., 2017) | https://emcore.ucsf.edu/ucsf-software |
| UCSF Chimera | (Pettersen et al., 2004) | https://www.cgl.ucsf.edu/chimera/ |
| Chimera X | (Goddard et al., 2018) | https://www.rbvi.ucsf.edu/chimerax/ |
| Coot | (Emsley and Cowtan, 2004) | https://www2.mrc-lmb.cam.ac.uk/personal/pemsley/coot/ |
| Cuemol2 | N/A | http://www.cuemol.org/ja/index.php?cuemol2 |
| Servalcat | (Yamashita et al., 2021) | https://github.com/keitaroyam/servalcat |
| GraphPad Prism | GraphPad | Graphpad.com |
| MATLAB | Mathworks | Mathworks.com |
| pClamp 10.6 | Molecular Devices | https://www.moleculardevices.com |
| Other |  |  |
| MultiClamp700B Amplifier | Molecular Devices | https://www.moleculardevices.com |
| DigiData 1440A | Molecular Devices | https://www.moleculardevices.com |
| DM-LFSA | Leica | N/A |
| Power meter | Thorlabs | PM100D |
| SPECTRA-X Light Engine | Lumencor | https://lumencor.com |
| R 1.2/1.3 grid: Au 300 mesh | QUANTIFOIL | Cat# N1-C14nAu30-01 |
| AMICON MWCO 10,000 | Merck / Millipore | Cat# UFC801024 |
| AMICON MWCO 50,000 | Merck / Millipore | Cat# UFC805024 |
| AKTA pure 25 L1 | Cytiva | Cat# 29018225 |
| Fluorescence detection HPLC system | Shimadzu | Model Prominence |
| V-750 UV-Visible Spectrophotometer | JASCO | https://jascoinc.com/products/spectroscopy/ |
| Vitrobot Mark IV | FEI / Thermo Fisher Scientific | https://www.thermofisher.com/jp/ja/home/electron-microscopy/products/sample-preparation-equipment-em/vitrobot-system.html |
| 300 kV Titan Krios microscope | FEI / Thermo Fisher Scientific | Out of production; https://www.thermofisher.com/jp/ja/home/electron-microscopy/products/transmission-electron-microscopes.html |
| K3 Summit camera with post-column energy filter | Gatan / Quantum | https://www.gatan.com/K3 |
| Innova S44i R | Eppendorf | Cat# S44I010226 |

### RESOURCE AVAILABILITY

#### Lead Contact

Further information and requests for resources and reagents should be directed to and will be fulfilled by the Lead Contact, Karl Deisseroth.

#### Material Availability

Plasmids/viruses and antibodies detailed in this manuscript are freely available for academic use.

#### Data and Code Availability

The raw images of *Hc*KCR1 and *Hc*KCR2 after motion correction has been deposited in the Electron Microscopy Public Image Archive, under accession EMPIAR-xxxxx and EMPIAR-xxxxx, respectively. The cryo-EM density map and atomic coordinates for *Hc*KCR1 and *Hc*KCR2 have been deposited in the Electron Microscopy Data Bank and PDB, under accessions EMD-xxxxx and xxxx, and EMD-xxxxx and xxxx, respectively. All other data are available upon request to the corresponding authors.

### EXPERIMENTAL MODEL AND SUBJECT DETAILS

#### Insect cell culture

*Spodoptera frugiperda (Sf9)* cells (Expression systems, authenticated by the vendor) were cultured in ESF921 medium (Expression systems) at 27.5°C with 130 rpm in an InnovaS44i R shaking incubator (Eppendorf).

#### HEK293 cell culture

HEK293FT cells (Thermo Fisher, authenticated by the vendor) were maintained in a 5% CO_2_ humid incubator with DMEM media (GIBCO) supplemented with 10% FBS (Invitrogen), and 1% Penicillin-Streptomycin (Invitrogen), and were enzymatically passaged at 90% confluence by trypsinization.

### METHOD DETAILS

#### Cloning, Protein Expression, and Purification of *Hc*KCR1 and *Hc*KCR2

Wild-type *Hc*KCR1 (M1-S265) was modified to include an N-terminal influenza hemagglutinin (HA) signal sequence and FLAG-tag epitope, and C-terminal enhanced green fluorescent protein (eGFP), followed by 10 × histidine and Rho1D4 epitope tags; the N-terminal and C-terminal tags are removable by human rhinovirus 3C protease cleavage. Wild type *Hc*KCR2 (M1-D265) was modified to include C-terminal Kir2.1 membrane targeting sequence, human rhinovirus 3C protease cleavage sequence, enhanced green fluorescent protein (eGFP), and 8 × histidine tag.

The constructs were expressed in *Spodoptera frugiperda* (*Sf9*) insect cells using the pFastBac baculovirus system. *Sf9* insect cells were grown in suspension to a density of 3.0 × 10^6^ cells ml^−1^, infected with baculovirus and shaken at 27.5°C for 24 h. All-*trans* retinal (ATR) (Sigma-Aldrich) is supplemented to a final concentration of 10 μM in the culture medium 24 h after the infection. The cell pellets were lysed with a hypotonic lysis buffer (20 mM HEPES-NaOH pH 7.5, 20 mM NaCl, 10 mM MgCl_2_, 1 mM benzamidine, 1 μg ml^−1^ leupeptin, 10 μM ATR), and cell pellets were collected by centrifugation at 10,000 ×g for 30 min. The above process was repeated twice; then, cell pellets were disrupted by homogenizing with a glass dounce homogenizer in a hypertonic lysis buffer (20 mM HEPES-NaOH pH 7.5, 1 M NaCl, 10 mM MgCl_2_, 1 mM benzamidine, 1 μg ml^−1^ leupeptin, 10 μM ATR), and crude membrane fraction was collected by ultracentrifugation (45Ti rotor, 125,000 ×g for 1 h). The above process was repeated twice; then, the membrane fraction was homogenized with a glass douncer in a membrane storage buffer (20 mM HEPES-NaOH pH 7.5, 500 mM NaCl, 10 mM imidazole, 20 % glycerol, 1 mM benzamidine, 1 μg ml^−1^ leupeptin), flash frozen in liquid nitrogen, and stored at −80°C until use.

The membrane fraction was solubilized in a solubilization buffer (1% *n*-dodecyl-β-D-maltoside (DDM) (EMD Millipore), 20 mM HEPES-NaOH pH 7.5, 500 mM NaCl, 20% glycerol, 5 mM imidazole, 1 mM benzamidine, 1 μg ml^−1^ leupeptin) and solubilized at 4°C for 2 h. The insoluble cell debris was removed by ultracentrifugation (45Ti rotor, 125,000 ×g, 1 h), and the supernatant was mixed with the Ni-NTA superflow resin (QIAGEN) at 4°C for 2 h. The Ni-NTA resin was loaded onto an open chromatography column, washed with 2.5 column volumes of wash buffer (0.05% DDM, 20 mM HEPES-NaOH pH7.5, 100 mM NaCl, and 25 mM imidazole) three times, and eluted by elution buffer (0.05% DDM, 20 mM HEPES-NaOH pH7.5, 100 mM NaCl, and 300 mM imidazole). After tag cleavage by His-tagged 3C protease, the sample was reapplied onto the Ni-NTA open column to trap the cleaved eGFP-His-tag and His-tagged 3C protease. The flow-through fraction was collected and concentrated to approximately 2 mg ml^−1^ using an Amicon ultra 50 kDa molecular weight cutoff centrifugal filter unit (Merck Millipore). The concentrated samples were ultracentrifuged (TLA 55 rotor, 71,680 ×g for 30 minutes) before size-exclusion chromatography on a Superdex 200 Increase 10/300 GL column (Cytiva), equilibrated in DDM SEC buffer (0.03% DDM, 20 mM HEPES-NaOH pH7.5, 100 mM NaCl). The peak fractions of the protein were collected and concentrated to approximately 10 mg ml^−1^..

#### Preparation of membrane scaffold protein

Membrane scaffold protein (MSP1D1E3) is expressed and purified as described earlier (Boldog et al., 2007) with the following modifications. Briefly, MSP1D1E3 gene in pET-43a(+) was transformed in *Escherichia coli* (*E. coli)* BL21 (DE3) cells. Cells were grown at 37°C with shaking to an OD_600_ of 0.5–1.0, and then expression of MSP1D1E3 was induced by addition of 1 mM IPTG. Cells were further grown for at 37°C 4 hr, and cells were harvested by centrifugation. Cell pellets were resuspended in PBS (–) buffer supplemented with 1% Triton X-100 and protease inhibitors and were lysed by sonication. The lysate was centrifuged at 30,000×*g* for 30 min, and the supernatant was loaded onto a Ni-NTA column equilibrated with lysis buffer, followed by washing with four bed volumes of wash buffer-1 (40 mM HEPES-NaOH pH7.5, 300 mM NaCl, 1% Triton X-100), four bed volumes of wash buffer-2 (40 mM HEPES-NaOH pH7.5, 300 mM NaCl, 50 mM sodium cholate), four bed volumes of wash buffer-3 (40 mM HEPES-NaOH pH7.5, 300 mM NaCl), four bed volumes of wash buffer-4 (40 mM HEPES-NaOH pH7.5, 300 mM NaCl, 20 mM imidazole), and eluted with wash buffer-4 containing 300 mM imidazole. The eluted MSP1D1E3 was dialyzed in buffer containing 10 mM HEPES-NaOH pH7.5, 100 mM NaCl, and concentrated to approximately 10 mg ml^−1^ using an Amicon ultra 10 kDa molecular weight cutoff centrifugal filter unit (Merck Millipore). The concentrated samples were ultracentrifuged (TLA 55 rotor, 71,680 ×g for 30 minutes), and stored at –80°C after flash freezing in liquid nitrogen. The concentration was determined by absorbance at 280 nm (extinction coefficient = 29,910 M^−1^ cm^−1^) measured by NanoDrop 2000c spectrophotometer (Thermo scientific).

#### Nanodisc reconstitution of *Hc*KCR1 and *Hc*KCR2

Prior to nanodisc reconstitution, 30 mg SoyPC (Sigma P3644-25G) was dissolved in 500 μL chloroform and dried using a nitrogen stream to form a lipid film. The residual chloroform was further removed by overnight vacuum desiccation. Lipid film were rehydrated in buffer containing 1% DDM, 20 mM HEPES-NaOH pH7.5, 100mM NaCl, resulting in a clear 10 mM lipid stock solution.

*Hc*KCR1 was reconstituted into nanodiscs formed by the scaffold protein MSP1E3D1 and SoyPC at a molar ratio of 1:4:400 (monomer ratio: HcKCR, MSP1E3D1, SoyPC). First, freshly purified *Hc*KCR1 in SEC buffer (0.05% DDM, 20 mM HEPES-NaOH pH7.5,100 mM NaCl) was mixed with SoyPC and incubated on ice for 20 min. Purified MSP1D1E3 was then added to mess up to total solution volume of 750 μL, and gently mixed on rotator at 4°C for 10 min. Final concentrations were 14.5 μM *Hc*KCR1, 58.2 μM MSP1E3D1, and 5.8 mM SoyPC, respectively. Detergents were removed by stepwise addition of Bio-Beads SM2 (Bio-Rad). Prior to use, Bio-Beads were washed by sonication in methanol, water, and buffer containing 20 mM HEPES-NaOH pH7.5,100 mM NaCl with an ultrasonic bath sonicator and weighed after liquid was removed by a P1000 tip. As the first batch, 100 mg Bio-Beads (final concentration of 133 mg ml^−1^) was added, and the mixture was gently rotated at 4°C for 12 h. The second batch of Bio-Beads (equal amount) was added and further rotated at 4°C for 2.5 h. The Bio-Beads were removed by passage through a PolyPrep column (Bio-Spin column, Bio-Rad), and the lysate was ultracentrifuged (TLA 55 rotor, 71,680 ×g for 30 minutes) before size-exclusion chromatography on a Superdex 200 Increase 10/300 GL column (Cytiva), equilibrated in buffer containing 20 mM HEPES-NaOH pH7.5, 100 mM NaCl. The peak fractions were collected and concentrated to approximately 6 mg ml^−1^ estimated based on the absorbance (A 280) value of 16, using an Amicon ultra 50 kDa molecular weight cutoff centrifugal filter unit (Merck Millipore).

*Hc*KCR2 was reconstituted into nanodiscs basically in the same manner as *Hc*KCR1. In brief, *Hc*KCR2, MSP1D1E3 and SoyPC were mixed at a molar ratio of 1:4:400, with the final concentration of 41 μM, 164 μM, and 4.1 mM, respectively. The total solution volume was 750 μL. Detergents were removed by stepwise addition of Bio-Beads SM2 (Bio-Rad). The first Bio-Beads batch amount was 25 mg. After rotation at 4°C for 12 h, 40 mg of fresh Bio-Beads were added every 12 h, twice in total. *Hc*KCR2 in a nanodisc was purified through size-exclusion chromatography and concentrated to approximately 12 mg ml^−1^ estimated based on the absorbance (A 280) value of 30, using an Amicon ultra 50 kDa molecular weight cutoff centrifugal filter unit (Merck Millipore).

#### Cryo-EM Grid Preparation of nanodisc-reconstituted *Hc*KCR1 and *Hc*KCR2

Prior to grid preparation, the sample was centrifuged at 20,380 ×g for 30 min at 4°C. The grids were glow-discharged with a PIB-10 plasma ion bombarder (Vacuum Device) at 10 mA current with the dial setting of 2 min for both side. 3 μL of protein solution was applied to freshly glow-discharged Quantifoil R1.2/1.3 Au 300 mesh holey carbon grid in dark room with dim red light. Samples were vitrified by plunging into liquid ethane cooled by liquid nitrogen with a FEI Vitrobot Mark IV (Thermo Fisher Scientific) at 4°C with 100% humidity. The blotting force was set as 10. The waiting and blotting time were 10 s and 4 s, respectively.

#### Cryo-EM data acquisition and image processing of *Hc*KCR1

Cryo-EM images were acquired at 300 kV on a Krios G3i microscope (Thermo Fisher Scientific) equipped with a Gatan BioQuantum energy filter and a K3 direct detection camera in the electron counting mode. The movie dataset was collected in standard mode, using a nine-hole image shift strategy in the SerialEM software (Mastronarde, 2005), with a nominal defocus range of −0.8 to −1.6 μm. The 5,445 movies were acquired at a dose rate of 14.313 e^-^/pixel/s, at a pixel size of 0.83 Å and a total dose of 48 e^-^/Å^2^.

The data processing was performed using the cryoSPARC v3.2.0 software packages (Punjani et al., 2017). The collected 5,445 movies were subjected to patch motion correction and patch CTF refinement in cryoSPARC. Initial particles were picked from all micrographs using blob picker and were extracted using a box size of 280 pixels. 407,781 particles were selected after 2D classification from 2,439,182 particles. The following *ab-initio* reconstruction, heterogeneous refinement, and non-uniform refinement (Punjani et al., 2020) enable us to reconstruct the 2.92 Å map (C1 symmetry) with 130,130 particles. Further particles were picked by template picker and Topaz picker (Bepler et al., 2019) and subjected to 2D classification followed by heterogeneous refinement. Non-uniform refinement after removing of the duplicated particles enable us to obtain 2.60 Å map (C3 symmetry) with 917,464 particles. The following 2D classification, global CTF refinement (Zivanov et al., 2020), and non-uniform refinement yielded the final map at a global resolution of 2.58 Å.

#### Cryo-EM data acquisition and image processing of *Hc*KCR2

Cryo-EM images were acquired at 300 kV on a Krios G4 microscope (Thermo Fisher Scientific) equipped with a Gatan BioQuantum energy filter and a K3 direct detection camera in the electron counting mode. The movie dataset was collected in standard mode, using the fringe free imaging (FFI) and aberration-free image shift (AFIS) strategy in the EPU software (Thermo Fisher Scientific), with a nominal defocus range of −0.6 to −1.6 μm. The 7,718 movies were acquired at a dose rate of 17.5 e^-^/pixel/s, at a pixel size of 0.83 Å and a total dose of 51 e^-^/Å^2^.

The data processing was performed using the cryoSPARC v3.3.2 software packages. The collected 7,718 movies were subjected to patch motion correction and patch CTF refinement in cryoSPARC. Particles were picked from all micrographs by blob picker, template picker, and Topaz picker, resulted in 3,382,955 particles, 5,852,598 particles, and 2,844,575 particles, respectively. These particle subsets were subjected to 2D classification and subsequent heterogeneous refinement. The particles in the best classes were 508,364 particles for blob picker, 777,572 particles for template picker, and 519,445 particles for Topaz picker, respectively. After removal of duplicates, 1,243,623 particles were selected and subjected to non-uniform refinement, resulting in a 2.66 Å map. The additional heterogeneous refinement, non-uniform refinement, local motion correction (Rubinstein and Brubaker, 2015), and another non-uniform refinement along with defocus refinement and global CTF refinement yielded the final map at a global resolution of 2.53 Å.

#### Model building and refinement

An initial model of *Hc*KCR1 was formed by rigid body fitting of the predicted models of *Hc*KCR1, generated using locally installed AlphaFold2 (Jumper et al., 2021). This starting model was then subjected to iterative rounds of manual and automated refinement in Coot (Emsley and Cowtan, 2004) and Refmac5 (Murshudov et al., 2011)(in Servalcat pipeline (Yamashita et al., 2021), respectively. The Refmac5 refinement was performed with the constraint of C3 symmetry. The initial model for *Hc*KCR2 was the refined model of *Hc*KCR1.

The final model was visually inspected for general fit to the map, and geometry was further evaluated using Molprobity (Chen et al., 2010). The final refinement statistics is summarized in Table S1. All molecular graphics figures were prepared with UCSF Chimera (Pettersen et al., 2004), UCSF ChimeraX (Goddard et al., 2018), CueMol2 (http://www.cuemol.org) and PyMOL (Schrödinger and DeLano, 2020)).

#### Pore analysis

The ion-conducting pore pathways were calculated by the software HOLLOW 1.3 with a grid-spacing of 1.0 Å (Ho and Gruswitz, 2008).

#### Measurement of UV absorption spectra and pH titration

To investigate the pH dependence of the absorption spectrum of *Hc*KCR1 and *Hc*KCR2, 10 mg ml^−1^ purified protein solution was 100-fold diluted in buffer containing 0.05 % DDM, 100 mM NaCl, and 100 mM of either citric acid pH2.2, citric acid pH 3.0, sodium acetate pH 4.0, sodium citrate pH 5.0, sodium cacodylate pH 6.0, HEPES-NaOH pH7.0, Tris-HCl pH8.0, *N*-cyclohexyl-2-aminoethanesulfonic acid (CHES) pH 9.0, 3-(cyclohexylamino)-1-propanesulfonic acid (CAPS) pH 10.0, or CAPS pH 11.0. The StockOptions pH Buffer Kit (Hampton research) was used for buffer preparation except for CHES pH 9.0 (Nacalai). The absorption spectra were measured with a V-750 UV-visible spectrometer (JASCO) at room temperature.

#### Laser flash photolysis

For the laser flash photolysis spectroscopy, *Hc*KCR1 wildtype and D105N were reconstituted in azolectin (11145, Sigma-Aldrich, Merck, Germany) with a protein-to-lipid molar ratio of 1:50 in 100 mM KCl, 20 mM HEPES-KOH pH 7.5. OD of the proteo-liposome suspensions was adjusted to ∼0.8 (protein concentration ∼0.2–0.3 mg/mL) at the absorption maximum wavelengths. The laser flash photolysis measurement was conducted as previously described (Inoue et al., 2013). Nano-second pulses from an optical parametric oscillator (5.7 mJ/pulse cm^2^, basiScan, Spectra-Physics, CA) pumped by the third harmonics of Nd–YAG laser (*λ* = 355 nm, INDI40, Spectra-Physics, CA) were used for the excitation of *Hc*KCR1 wildtype and D105N at *λ*_exc_ = 510 and 500 nm, respectively. The transient absorption spectra were obtained by monitoring the intensity change of white-light from a Xe-arc lamp (L9289-01, Hamamatsu Photonics, Japan) passed through the sample with an ICCD linear array detector (C8808-01, Hamamatsu, Japan). To increase the signal-to-noise (S/N) ratio, 45–60 spectra were averaged, and the singular-value-decomposition (SVD) analysis was applied. To measure the time-evolution of transient absorption change at specific wavelengths, the output of a Xe-arc lamp (L9289-01, Hamamatsu Photonics, Japan) was monochromated by monochromators (S-10, SOMA OPTICS, Japan) and the change in the intensity after the photo-excitation was monitored with a photomultiplier tube (R10699, Hamamatsu Photonics, Japan). To increase the S/N ratio, 100–200 signals were averaged.

To measure the transient absorption change of pyranine due to proton release and uptake by *Hc*KCR1 wildtype, the protein was solubilized in 100 mM KCl, 0.05% DDM, and pH was adjusted to 7.2 close to the pKa of pyranine by adding NaOH, and then 40 µM pyranine (L11252, Wako, Japan) was added. The formation and disappearance of the protonated form of pyranine were monitored at 454 nm by subtracting the transient absorption change obtained without pyranine from that obtained with pyranine as previously reported (Inoue et al., 2018).

#### High performance liquid chromatography (HPLC) analysis of retinal isomers in *Hc*KCR1

The HPLC analysis of retinal isomers was conducted as described elsewhere (Kishi et al., 2022) with a slight modification. The purified sample was incubated at 4°C overnight in the dark prior to the HPLC analysis. A 30-μL sample and 120 μL of 90% (v/v) methanol aqueous solution and 10 μL of 2 M hydroxylamine (NH_2_OH) were added to the sample. Then, retinal oxime hydrolyzed from the retinal chromophore in *Hc*KCR1 was extracted with 500 μL of *n*-hexane. A 200 μL of the extract was injected into an HPLC system equipped with a silica column (particle size 3 μm, 150 × 6.0 mm; Pack SIL, YMC, Japan), a pump (PU-4580, JASCO, Japan), and a UV–visible detector (UV-4570, JASCO, Japan). As the mobile-phase solvent, *n*-hexane containing 15% ethyl acetate and 0.15 % ethanol was used at a flow rate of 1.0 mL min^−1^. Illumination was performed with green light (510 ± 5 nm) for 60 s. The molar composition of the retinal isomers the sample was calculated with the molar extinction coefficient at 360 nm for each isomer (all-*trans*-15-*syn*: 54,900 M^−1^ cm^−1^; all-*trans*-15-*anti*: 51,600 M^−1^ cm^−1^; 13-*cis*-15-*syn*, 49,000 M^−1^ cm^−1^; 13-*cis*-15-*anti*: 52,100 M^−1^ cm^−1^; 11-*cis*-15-*syn*: 35,000 M^−1^ cm^−1^; 11-*cis*-15-*anti*: 29,600 M^−1^ cm^−1^; 9-*cis*-15-*syn*: 39,300 M^−1^ cm^−1^; 9-*cis*-15-*anti*: 30,600 M^−1^ cm^−1^) (Groenendijk et al., 1979; Ozaki et al., 1986).

#### Laser patch clamp

The electrophysiological assays of *Hc*KCR1 were carried out using ND7/23 cells, as described previously (Nagasaka et al., 2020) with a slight modification. Briefly, ND7/23 cells were grown in Dulbecco’s modified Eagle’s medium (D-MEM, FUJIFILM Wako Pure Chemical Co., Osaka, Japan) supplemented with 5% fetal bovine serum (FBS) under a 5% CO_2_ atmosphere at 37°C. Eight hours after the transfection, the medium was replaced by D-MEM containing 5% FBS, 50 ng/mL nerve growth factor-7S (Sigma-Aldrich, St. Louis, MO), 1 mM N6,2’-O-dibutyryladenosine-3’,5’-cyclic monophosphate sodium salt (Nacalai tesque, Kyoto, Japan), and 1 μM Cytosine-1-β-D(+)-arabinofuranoside (FUJIFILM Wako Pure Chemical Co., Osaka, Japan). The coding sequence of *Hc*KCR1 was fused to a Kir2.1 membrane trafficking signal, eYFP, and an ER-export signal (Gradinaru et al., 2010). The gene was cloned into a vector behind a CMV-promotor and the expression plasmids were transiently transfected in ND7/23 cells using LipofectamineTM 3000 transfection reagent (Thermo Fisher Scientific Inc., Waltham, MA) and electrophysiological recordings were conducted at 2–3 days after the transfection. The transfected cells were identified by the presence of eYFP fluorescence under an up-right microscope (BX50WI, Olympus, Tokyo, Japan).

All experiments were carried out at room temperature (20–22 °C). Currents were recorded using an EPC-8 amplifier (HEKA Electronic, Lambrecht, Germany) under a whole-cell patch clamp configuration. The internal pipette solution contained 121.2 mM KOH, 90.9 mM glutamate, 5 mM Na_2_EGTA, 49.2 mM HEPES, 2.53 mM MgCl_2_, 2.5 mM MgATP, 0.0025 mM ATR (pH 7.4 adjusted with HCl). Extracellular solution contained 138 mM NaCl, 3 mM KCl, 2.5 mM CaCl_2_, 1 mM MgCl_2_, 4 mM NaOH, and 10 mM HEPES at pH 7.4 (with 11 mM glucose added up to 310 mOsm). The pipette resistance was adjusted to 3–6 MΩ (3.7 ± 0.4, n = 7) with a series resistance of 6–11 MΩ (8.3 ± 0.8) and a cell capacitance of 32–216 pF (83 ± 21) with the extracellular/intracellular solutions. In every experiment, the series resistance was compensated. While voltage-clamping at a holding potential, a laser flash (3–5 ns) at 532 nm (Nd:YAG laser, Minilite II, Continuum, San Jose, CA) was illuminated through an objective lens (LUMPlan FL 40x, NA 0.80W, Olympus, Japan). The timing of laser flash was set to be time 0 according to the photodiode response under the sample. The measurements were conducted with a holding potential of 0 mV at every 15 s. The data were filtered at 1 kHz, sampled at 250 kHz (Digidata1440 A/D, Molecular Devices Co., Sunnyvale, CA), collected using pClamp10.3 software (Molecular Devices Co., Sunnyvale, CA), and stored in a computer. Five current responses were averaged and served for the following analyses. Using the simplex method of nonlinear least-squares (IgorPro 9, WaveMetrics, Portland, OR), the kinetics of photocurrent were fitted by a triple-exponential function.

#### ATR-FTIR Spectroscopy

Ion binding to *Hc*KCR1 was monitored by ATR-FTIR spectroscopy as described previously (1-5)(Furutani et al., 2011; Hashimoto et al., 2020; Inoue et al., 2013; Iwaki et al., 2018; Katayama et al., 2018), except for some minor modifications for reconstitution into the membrane. In ATR-FTIR spectroscopy, rhodopsins are normally reconstituted into lipids for forming a film on the ATR-prism. Thus, sample was reconstituted with a protein-to-lipid (asolectin; Sigma-Aldrich) molar ratio of 1:20, by removing the n-dodecyl-β-D-maltoside (DDM) with Bio-Beads (SM-2, Bio-Rad) at 4 °C in dark condition. The *Hc*KCR1 sample in asolectin liposomes was washed repeatedly with a buffer containing 2 mM K_2_HPO_4_ / KH_2_ PO_4_ (pH 7.5) and collected by ultracentrifuging for 20 min at 222,000 x g at 4 °C in dark condition. The lipid-reconstituted *Hc*KCR1 was placed on the surface of a silicon ATR crystal (Smiths, three internal total reflections) and naturally dried. The sample was then rehydrated with the buffer at a flow rate of 0.6 ml min^-1^, and temperature was maintained at 20 °C by circulating water. The perfusion buffer is composed of 200 mM NaCl, 200 mM Tris-HCl, pH 7.5 (buffer A) and 200 mM KCl, 200 mM Tris-HCl, pH 7.5 (buffer B). In the case of anion binding experiments, the perfusion buffer was replaced with 200 mM NaCl, 20 mM HEPES-NaOH, pH 7.5 (buffer A) and 200 mM NaBr, 20 mM HEPES-NaOH, pH 7.5 (buffer B), respectively.

ATR-FTIR spectra were recorded in kinetics mode at 2 cm^−1^ resolution, renge of 4000-700 cm^−1^ using an FTIR spectrometer (Agilent) equipped with a liquid nitrogen-cooled mercury-cadmium-telluride (MCT) detector (an average of 1710 interferograms per 15 min). Ion binding-induced difference spectra were measured by exchanging the buffer A and buffer B. The cycling procedure is shown in Figure S3G, and the difference spectra were calculated as the averaged spectra in buffer B minus buffer A. The spectral contributions of the unbound salt, the protein-lipid swelling/shrinkage, and the water-buffer components were corrected as described previously (Hashimoto et al., 2020).

Light-induced structural changes of *Hc*KCR1 were also measured by ATR–FTIR as shown in Figure S3G. Since ATR-FTIR experimental setup has been optimized for ion perfusion-induced difference spectroscopy using a solution exchange system, we have modified experimental setup that enables light irradiation experiment. A light source was installed above the ATR prism. In addition, an optical filter and a condenser lens were placed directly under the light source. To obtain the ion binding-induced difference spectra under the light illumination condition, light minus dark difference spectra under perfusing the different solution between buffer A and buffer B was subtracted each other. The spectral contributions of the unbound salt, the protein-lipid swelling/shrinkage, and the water-buffer components were also corrected as described previously (Hashimoto et al., 2020).

#### *In vitro* electrophysiology

Cells and devices for the measurement were prepared as described (Kishi et al., 2022). Briefly, HEK293 cells (Thermo Fisher) expressing opsins were placed in an extracellular tyrode medium (150 mM NaCl, 4 mM KCl, 2 mM CaCl_2_, 2 mM MgCl_2_, 10 mM HEPES pH 7.4, and 10 mM glucose). Borosilicate pipettes (Harvard Apparatus, with resistance of 4 – 6 MOhm) were filled with intracellular medium (140 mM potassium-gluconate, 10 mM EGTA, 2 mM MgCl_2_ and 10 mM HEPES pH 7.2). Light was delivered with the Lumencor Spectra X Light engine with 470 nm and 560 nm filters for blue and orange light delivery, respectively.

Channel kinetics and photocurrent amplitudes were measured in voltage clamp mode at 0 mV (before liquid junction potential correction) holding potential and then analyzed in Clampfit software (Axon Instruments) after smoothening using a lowpass Gaussian filter with a −3 dB cutoff for signal attenuation and noise reduction at 1,000 Hz. Liquid junction potentials were corrected using the Clampex built-in liquid junction potential calculator as previously described (Kishi et al., 2022). Equilibrium potentials were measured by holding membrane potentials from −96 mV (after LJP correction) in steps of 10 mV.

Statistical analysis was performed with one-way ANOVA and the Kruskal–Wallis test for non-parametric data, using Prism 7 (GraphPad) software. Data collection across opsins was randomized and distributed to minimize across-group differences in expression time, room temperature, and related experimental factors.

#### Ion selectivity testing in HEK293 cells

HEK293 cells and devices for the measurement were prepared as described in the previous section. For the high sodium extracellular / high potassium intracellular condition, we used sodium bath solution containing 150 mM NaCl, 2 mM CaCl_2_, 2 mM MgCl_2_, 10 mM HEPES pH 7.3 with 10 mM glucose, along with potassium pipette solution containing 150 mM KCl, 2 mM CaCl_2_, 2 mM MgCl_2_, 10mM HEPES pH 7.2, and 10 mM glucose. For the high potassium extracellular / high sodium intracellular condition, NaCl and KCl concentrations were reversed, and all other ionic concentrations were kept constant. Liquid junction potentials were corrected using the Clampex built-in liquid junction potential calculator.

For sodium extracellular/potassium intracellular condition, equilibrium potentials were measured by holding membrane potentials from −124 mV to + 16 mV in steps of 10 mV (after LJP correction). for potassium extracellular/sodium intracellular, equilibrium potentials were measured by holding from −26 mV to 104 mV (after LJP correction) in 10 mV steps. The relative ion permeability of sodium and potassium (*P*_K_/*P*_Na_) was calculated using the Goldman-Hodgkin-Katz equation, with T = 298K, *P_Cl_* = 0, F = 96485 C/mol and R = 8.314 J/K·mol.

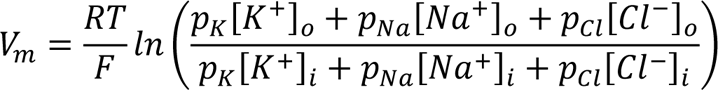

To test intracellular Guanidinium blockage, an intracellular buffer containing 150 mM GuHCl, 2 mM CaCl_2_, 2 mM MgCl_2_, 10 mM HEPES pH 7.3, and 10 mM glucose was used with a regular high sodium extracellular buffer.

Statistical analysis was performed with one-way ANOVA and the Kruskal–Wallis test for non-parametric data, using Prism 7 (GraphPad) software. Data collection across opsins was randomized and distributed to minimize across-group differences in expression time, room temperature, and related experimental factors.

#### System setup for molecular dynamics simulations

We performed simulations of *Hc*KCR1 WT and a *Hc*KCR1 D116N mutant. The simulations were initiated using structure reported in this manuscript. For the D116N mutant simulations, the mutation was introduced while maintaining the positions of all the common atoms. To study the exit pathway of ions from the channel, we placed a K^+^ in both the intracellular and extracellular vestibules near D166 and D205, respectively. For each simulation condition, we performed three independent simulations, each 500 ns in length. For each simulation, initial atom velocities were assigned randomly and independently.

The structure was aligned to the Orientations of Proteins in Membranes(Lomize et al., 2006) entry for 1M0L(Schobert et al., 2002) (bacteriorhodopsin). Prime (Schrödinger)(Jacobson et al., 2002) was used to model missing side chains, and to add capping groups to protein chain termini. The Crosslink Proteins tool (Schrödinger) was used to model unresolved portions of ECL2, ICL3, and ECL3. Parameters for the ligands were generated using the Paramchem webserver(Vanommeslaeghe and MacKerell, 2012; Vanommeslaeghe et al., 2010, 2012). Dowser software was used to add waters to cavities within the protein structure(Zhang and Hermans, 1996). Six POPC lipids were modeled in the center of the trimer; three in the extracellular leaflet and three in the intracellular leaflet. Protonation states of all titratable residues were assigned at pH 7. Histidine residues were modeled as neutral, with a hydrogen atom bound to either the delta or epsilon nitrogen depending on which tautomeric state optimized the local hydrogen-bonding network. Using Dabble(Betz, 2017), the prepared protein structures were inserted into a pre-equilibrated palmitoyl-oleoyl-phosphatidylcholine (POPC) bilayer, the system was solvated, and potassium and chloride ions were added to neutralize the system and to obtain a final concentration of 150 mM. The final systems comprised approximately 101,000 atoms, and system dimensions were approximately 105×105×95 Å.

#### Molecular dynamics simulation and analysis protocols

We used the CHARMM36m force field for proteins, the CHARMM36 force field for lipids and ions, and the TIP3P model for waters(Guvench et al., 2008; Huang et al., 2017; Klauda et al., 2010). Retinal parameters were obtained through personal communication with Scott Feller(Zhu et al., 2013). All simulations were performed using the Compute Unified Device Architecture (CUDA) version of particle-mesh Ewald molecular dynamics (PMEMD) in AMBER18(Lee et al., 2018) on graphics processing units (GPUs).

Systems were first minimized using three rounds of minimization, each consisting of 500 cycles of steepest descent followed by 500 cycles of conjugate gradient optimization. 10.0 and 5.0 kcal·mol^−1^·Å^−2^ harmonic restraints were applied to protein, lipids, and ligand for the first and second rounds of minimization, respectively. 1 kcal·mol^−1^·Å^−2^ harmonic restraints were applied to protein and ligand for the third round of minimization. Systems were then heated from 0 K to 100 K in the NVT ensemble over 12.5 ps and then from 100 K to 298 K in the NPT ensemble over 125 ps, using 10.0 kcal·mol^−1^·Å^−2^ harmonic restraints applied to protein and ligand heavy atoms. Subsequently, systems were equilibrated at 298 K and 1 bar in the NPT ensemble, with harmonic restraints on the protein and ligand non-hydrogen atoms tapered off by 1.0 kcal·mol^-1^·Å^-2^ starting at 5.0 kcal·mol^-1^·Å^-2^ in a stepwise fashion every 2 ns for 10 ns, and then by 0.1 kcal·mol^-1^·Å^-2^ every 2 ns for 20 ns. Production simulations were performed without restraints at 310 K and 1 bar in the NPT ensemble using the Langevin thermostat and the Monte Carlo barostat, and using a timestep of 4.0 fs with hydrogen mass repartitioning(Hopkins et al., 2015). Bond lengths were constrained using the SHAKE algorithm(Ryckaert et al., 1977). Non-bonded interactions were cut off at 9.0 Å, and long-range electrostatic interactions were calculated using the particle-mesh Ewald (PME) method with an Ewald coefficient of approximately 0.31 Å, and 4th order B-splines. The PME grid size was chosen such that the width of a grid cell was approximately 1 Å. Trajectory frames were saved every 200 ps during the production simulations.

### QUANTIFICATION AND STATISTICAL ANALYSIS

Statistical analysis was performed with one-way ANOVA and the Kruskal–Wallis test for non-parametric data, using Prism 7 (GraphPad) software. Data collection across opsins was randomized and distributed to minimize across-group differences in expression time, room temperature, and related experimental factors.

